# Dectin-1 Stimulation Promotes a Distinct Inflammatory Signature in the Setting of HIV-infection and Aging

**DOI:** 10.1101/2022.09.01.505710

**Authors:** Archit Kumar, Jiawei Wang, Haowen Zhou, Chris Radcliffe, Brent Vander Wyk, Heather G. Allore, Sui Tsang, Lydia Barakat, Subhasis Mohanty, Hongyu Zhao, Albert C. Shaw, Heidi J. Zapata

**Author notes:** Corresponding Author: Heidi J Zapata: PO Box 208022, 300 Cedar Street, New Haven, CT, 06520-8022.

## Abstract

Dectin-1 is an innate immune receptor that recognizes and binds β-1,3/1,6 glucans on fungi. We evaluated Dectin-1 function in myeloid cells in a cohort of HIV-positive and HIV-negative young and older adults. Stimulation of monocytes with β-D-glucans induced a pro-inflammatory phenotype in monocytes of HIV-infected individuals that was characterized by increased levels of IL-12, TNF-α, and IL-6, with some age-associated cytokine increases also noted. Dendritic cells showed a striking HIV-associated increase in IFN-α production. These increases in cytokine production paralleled increases in Dectin-1 surface expression in both monocytes and dendritic cells that were noted with both HIV and aging. Differential gene expression analysis showed that HIV-positive older adults had a distinct gene signature compared to other cohorts characterized by a robust TNF-α and coagulation response (increased at baseline), a persistent IFN-α and IFN-γ response, and an activated dendritic cell signature/M1 macrophage signature upon Dectin-1 stimulation. Dectin-1 stimulation induced a strong upregulation of MTORC1 signaling in all cohorts, although increased in the HIV-Older cohort (stimulation and baseline). Overall, our study demonstrates that the HIV Aging population has a distinct immune signature in response to Dectin-1 stimulation. This signature may contribute to the pro-inflammatory environment that is associated with HIV and Aging.

## Introduction

Dectin-1(CLEC7A) is a member of the C-type lectin receptor family that recognizes carbohydrate structures on pathogens, with Dectin-1 specifically recognizing and binding β-1,3/1,6--glucans (Drummond & Brown, 2011; Plato, Willment, & Brown, 2013; Taylor et al., 2007), a component of the fungal cell wall. Dectin-1, originally identified as “dendritic-cell-associated C-type lectin-1” (Yokota, Takashima, Bergstresser, & Ariizumi, 2001) is primarily expressed on myeloid cells, including dendritic cells, monocytes, macrophages, and neutrophils. Dectin-1 expression has also been noted on non-immune cells such as lung epithelium (Heyl et al., 2014) and intestinal M cells (Rochereau et al., 2013). Through the recognition of β-glucans, Dectin-1 can recognize human pathogens that are particularly important causes for infection in immunocompromised hosts, such as *Aspergillus, Candida, Coccidioides, Talaromyces*, and *Pneumocystis* (Drummond & Brown, 2011; Plato et al., 2013). Binding and stimulation of Dectin-1 leads to phosphorylation of its ITAM motif, subsequent downstream signaling through the Syk/CARD9 pathway that ultimately results in NF-κB activation and inflammatory cytokine production. Dectin-1 activation also induces activation of the NLRP3 inflammasome and a Th-17 response (Plato et al., 2013)—playing a major role in anti-fungal immunity. Additionally, Dectin-1 may also recognize other microbial pathogens such as bacteria (*Salmonella*),mycobacteria (Kalia, Singh, & Kaur, 2021; Wagener, Hoving, Ndlovu, & Marakalala, 2018) as well as endogenous ligands (Deerhake & Shinohara, 2021) that may contribute to age-related chronic inflammation.

Invasive fungal infections in humans carry unacceptably high rates of mortality that can exceed 50% despite anti-fungal therapies. Patients with HIV-infection (Brown et al., 2012), and increased age (Antiretroviral Therapy Cohort, 2008; Samji et al., 2013) are at high risk but the effects of HIV-infection and age on Dectin-1 function remain unknown. Consequently, we evaluated Dectin-1 function in a cohort of HIV-infected and uninfected young and older adults. Peripheral mononuclear cells (PBMCs) were stimulated with whole glucan particles (1,3/1,6-β-glucan from *S. cerevisiae*) which specifically stimulate Dectin-1 and lack TLR stimulating activity. Using multicolor flow cytometry, we evaluated monocytes, the most abundant innate cells in the peripheral blood functioning as antigen presenting cells that may differentiate into macrophages and dendritic cells at sites of inflammation. Monocytes can be subdivided into the following subsets as defined by their expression of CD14 and CD16: classical (CD14+CD16lo), inflammatory (CD14+CD16+, known to be expanded in the setting of aging and HIV infection)(Zapata & Shaw, 2014a), and non-classical (CD14lo CD16hi) subset (also increased with HIV disease). Monocytes can also be further defined by the expression of activation markers such as CD11b of which the expression is also increased in the setting of aging and HIV infection (Dutertre et al., 2012; Metcalf et al., 2017; Shaw, Goldstein, & Montgomery, 2013; Zapata & Shaw, 2014b; Ziegler-Heitbrock & Hofer, 2013). Dectin-1 function was also studied in human dendritic cells which are potent antigen presenting cells found in the peripheral blood and lymphoid tissue. Dendritic cells can be subdivided into myeloid dendritic cells (CD11c+, CD123-; mDCs), and plasmacytoid dendritic cells (CD11c-, CD123+; pDCs) via expression of CD11c and CD123 (Panda et al., 2010) as previously described. Finally, we performed RNA-seq analysis on sorted inflammatory monocytes to elucidate the effects of age and HIV infection on transcriptomic signatures at baseline and upon Dectin-1 activation.

## Results

We recruited a total of 81 HIV-negative and HIV-positive participants, divided between younger and older adults (as noted in the methods); the characteristics of enrolled individuals are shown in Table 1. The HIV-positive and HIV-negative groups were comparable in age and gender distribution, incidence of comorbidities such as diabetes, metabolic syndrome, cardiovascular disease and pulmonary disease. HIV-positive adults, compared to the HIV-negative cohort, had higher rates of recreational drug use, number of co-morbidities and differed in distribution of self-reported race and Hispanic ethnicity. Nearly all of the HIV-infected cohort were on antiretroviral therapy (ART) and had CD4 counts >200/mm^3^. However, to account for the effects of differing years with HIV-infection, percent lifespan with HIV-infection (since diagnosis) was calculated for each HIV-infected subject and is represented as a median with an interquartile range (25 percentile, 75 percentile). Of note, there were four congenitally HIV-infected subjects, all in the young group.

**Table 1.** Descriptive Statistics stratified by HIV Status (N=81). Demographic information for HIV-positive and HIV-negative cohorts. Metabolic syndrome = DM + HTN + HLD, PPD= purified protein derivative. All values are n (%). Unless otherwise noted, significance was evaluated using a chi-square test. P values less than or equal to 0.05 were considered significant. The age groups include younger = 21-35, and (≥ 60 years) for most experiments (Monocytes). However, the age groups were extended to 21-40 and (≥ 50 years) for the Dendritic cell experiments. ^1^ Significance assessed using Fisher’s Exact Test ^2^ The value of % life span with HIV (since diagnosis) shown in the table represents an interquartile range, with the median (25 percentile, 75 percentile).

|  |  | Monocytes |  | p-value | Dendritic cells |  | p-value |
| --- | --- | --- | --- | --- | --- | --- | --- |
|  |  | HIV Positive | HIV Negative |  | HIV Positive | HIV Negative |  |
| **Total** |  | 36 | 45 |  | 27 | 39 |  |
| **Age Group** | Younger | 14 (38.9) | 21 (46.7) | 0.483 | 10 (37.0) | 21 (53.9) | 0.215 |
|  | Older | 22 (61.1) | 24 (53.3) |  | 17 (63.0) | 18 (46.2) |  |
| **Gender** | Male | 21 (58.3) | 20 (44.4) | 0.214 | 16 (59.3) | 19 (48.7) | 0.399 |
|  | Female | 15 (41.7) | 25 (55.6) |  | 11 (40.7) | 20 (51.3) |  |
| **Race/Ethnicity** | Hispanic | 10 (27.8) | 3 (6.7) | 0.014<sup>1</sup> | 5 (18.5) | 1 (2.6) | 0.038<sup>1</sup> |
|  | White | 14 (38.9) | 33 (73.3) | 0.001<sup>1</sup> | 13 (48.1) | 32 (82.1) | <.001<sup>1</sup> |
|  | Black | 15 (41.7) | 6 (13.3) |  | 9 (33.3) | 3 (7.7) |  |
|  | Asian | 0 (0) | 4 (8.9) |  | 0 (0) | 4 (10.3) |  |
|  | Other | 7 (19.4) | 2 (4.4) |  | 5 (18.5) | 0 (0) |  |
| **Smoking** |  | 9 (25) | 5 (11.1) | 0.14<sup>1</sup> | 7 (25.9) | 3 (7.7) | 0.077<sup>1</sup> |
| **Recreational Drugs** |  | 11 (30.6) | 2 (4.4) | 0.002<sup>1</sup> | 9 (33.3) | 1 (2.6) | <.001<sup>1</sup> |
| **Number of Comorbidities** | 0 | 0 (0) | 6 (13.3) | <.001<sup>1</sup> | 0 (0) | 6 (15.4) | .024<sup>1</sup> |
|  | 1-3 | 10 (27.8) | 17 (37.8) |  | 7 (25.9) | 17 (43.6) |  |
|  | 4-7 | 19 (52.8) | 12 (26.7) |  | 11 (40.7) | 11 (28.2) |  |
|  | >7 | 7 (19.4) | 10 (22.2) |  | 9 (33.3) | 5 (12.8) |  |
| **History of positive PPD / Quantiferon** |  | 6 (16.7) | 2 (4.5) | 0.131<sup>1</sup> | 4 (14.8) | 2 (5.3) | 0.224<sup>1</sup> |
| **Diabetes** |  | 8 (22.2) | 11 (24.4) | 0.815 | 6 (22.2) | 7 (17.9) | 0.458 |
| **Metabolic Syndrome** |  | 5 (13.9) | 11 (24.4) | 0.273<sup>1</sup> | 4 (14.8) | 7 (17.9) | 1.000 |
| **Cardiovascular Disease** |  | 10 (27.8) | 12 (26.7) | 0.911 | 10 (37) | 9 (23.1) | 0.218 |
| **Pulmonary Disease** |  | 11 (30.6) | 12 (26.7) | 0.700 | 7 (25.9) | 9 (23.1) | 0.791 |
| **CD4 count >200** |  | 35 (97.2) |  |  | 26 (96.3) |  |  |
| **On ART** |  | 34 (94.4) |  |  | 25 (92.6) |  |  |
| **HIV VL > 100** |  | 4 (11.1) |  |  | 7 (25.9) | 2 (5.1) |  |
| **% life span with HIV** |  | 26.1 (18.4-44.6)<sup>2</sup> |  |  | 34.3 (12.1-49.1)<sup>2</sup> |  |  |

Freshly isolated PBMCs were stimulated with whole glucan particles (1,3/1,6-β-glucan, WGP Dispersable), a specific Dectin-1 agonist (Goodridge et al., 2011) and analyzed via intracellular cytokine staining (ICS) and multicolor flow cytometry. A soluble form of WGP that binds to the Dectin-1 receptor without activation was used as a negative control as noted in the methods. The soluble form of WGP did not elicit any cytokine production (data not shown). We determined intracellular IL-10, IL-12, IL-6 and TNF-α production in classical (CD14+CD16lo), inflammatory (CD14+CD16+), activated (CD11b+CD14+) and non-classical monocytes (CD14loCD16H) monocytes (Ziegler-Heitbrock & Hofer, 2013). In activated CD11b+ CD14+ monocytes, we found an age-associated increase in Dectin-1-induced IL-6, IL-12 and TNF-α production in older, compared to young, HIV-negative adults. Furthermore, TNF-α and IL-12 production in these activated monocytes was increased in both young and older adults with HIV infection, when compared to HIV-negative adults, respectively (Fig. 1a). Robust IL-10 production appeared comparable in all cohorts. These results in CD11b+ monocytes reflect an age- and HIV-associated heightened activation of Dectin-1 function. CD14+ CD16+ inflammatory monocytes also showed an HIV-associated increase in TNF-α production (HIV-positive vs. HIV negative individuals) in both young and older adults. An HIV-associated increase in IL-12 levels was also noted in HIV-positive young, when compared to HIV-negative young individuals (Figure 1b). Classical monocytes (CD14+CD16lo) demonstrated an age-associated increase in production of IL-12 and IL-6, but in contrast to CD14+ CD11b+ or inflammatory monocytes a statistically significant effect of HIV infection was not found (Supplemental Figure 2a). In contrast, non-classical CD14+CD16H monocytes from HIV-negative adults showed a significant age-associated decrease in intracellular IL-12 levels, with a trend of decreased TNF-α production that did not reach significance. Cytokine production of IL-12 and TNF-α in HIV-positive young and older adults was comparable to the young HIV-negative cohort (Supplemental Figure 2 a, b). Taken together, these results indicate that activated CD11b+ and inflammatory monocytes show a HIV-associated increase in TNF-α and IL-12 production. In the CD11b+ activated subset, we also observed an age associated increase in IL-12, TNF-α and IL-6 production.

**Figure 1.**
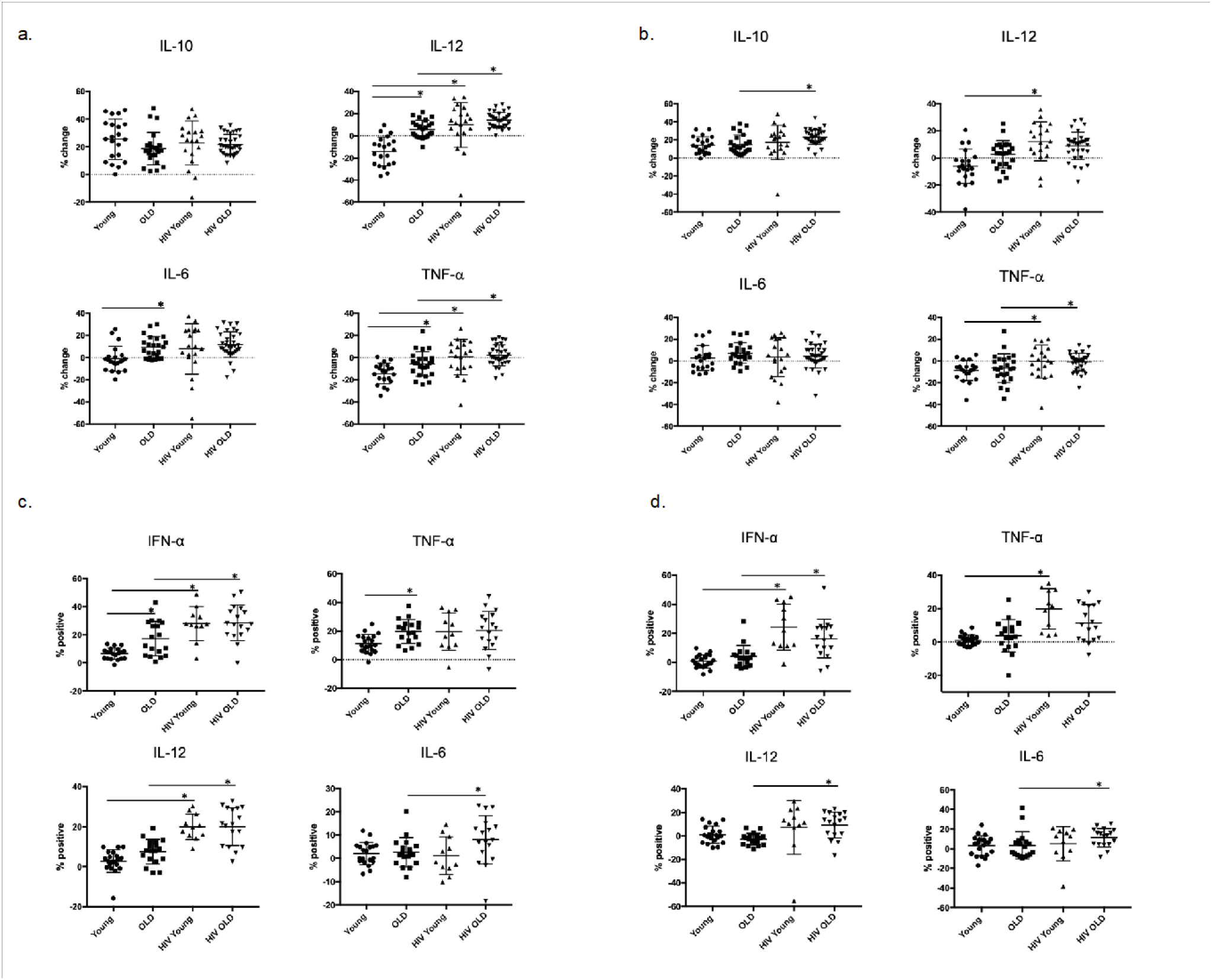
Effects of age and HIV-infection on Dectin-1 induced cytokine production in Monocytes and Dendritic cells. HIV-negative young adults (n=21), HIV-negative older adults (n= 24), HIV-positive young adults (n =14), and HIV-positive older adults (n = 22). Dot plots showing percent change in production of interleukin IL-10, IL-12 (p70 isoform), IL-6, and tumor necrosis factor-α (TNF-α) compared to baseline after stimulation with whole glucan particles (WGP or 1,3/1,6-β-glucan). The following comparisons indicated by asterisks were statistically significant using a Wilcoxon two-sample test with *t* approximation, which were then adjusted with a false discover rate (FDR) calculation for multiple comparisons. **a) Total cytokine production in CD11b+ Activated monocytes**. HIV-negative older adults versus young adults: IL-12 (p= 0.00065), IL-6 (p= 0.006), TNF-α (p= 0.026), HIV-positive young adults versus HIV-negative young adults: IL-12 (p=0.011), TNF-α (p=0.016), HIV-positive older adults versus HIV-negative older adults: IL-12 (p=0.01), TNF-α (0.0057). All other comparisons were not significant. **b) Total cytokine production in Inflammatory (CD14+CD16+) monocytes**. HIV-positive young adults versus HIV-negative young adults: IL-12 (p=0.0199), TNF-α (p=0.024), HIV-positive older adults versus HIV-negative older adults: IL-10 (p= 0.011), TNF-α (0.032). All other comparisons were not significant. c) **Total Cytokine Production in Myeloid Dendritic Cells**. HIV-negative young adults (n=21), HIV-negative older adults (n= 18), HIV-positive young adults (n =10), and HIV-positive older adults (n = 17). Dot plots showing percent change in production of interleukin IFN-α, TNF-α, IL-12, and IL-6 compared to baseline after stimulation with whole glucan particles (WGP or 1,3/1,6-β-glucan) (labeled as percent positive). The following comparisons indicated by asterisks were statistically significant using a Wilcoxon two-sample test with *t* approximation, which were then adjusted with a false discovery rate (FDR) calculation for multiple comparisons. **i) IFN-α**, HIV-negative older adults vs. HIV-negative young adults (p=0.0373), HIV-positive young adults vs. HIV-negative young adults (p=0.0004), HIV-positive older adults vs. HIV-negative older adults (p=0.011). All other comparisons were not significant. **ii) TNF**-**α**, HIV-negative older adults vs. HIV-negative young adults (p=0.023), All other comparisons were not significant. **iii) IL-12**, HIV-positive young adults vs. HIV-negative young adults (p=0.0001), HIV-positive older adults vs. HIV-negative older adults (p=0.001) **iiii) IL-6**, HIV-positive older adults vs. HIV-negative older adults (p=0.035). d) **Total Cytokine Production in Plasmacytoid Dendritic Cells**. HIV-negative young adults (n=21), HIV-negative older adults (n= 18), HIV-positive young adults (n =10), and HIV-positive older adults (n = 17). Dot plots showing percent change in production of interleukin IFN-α, TNF-α, IL-12, and IL-6 compared to baseline after stimulation with whole glucan particles (WGP or 1,3/1,6-β-glucan) (labeled as percent positive). The following comparisons indicated by asterisks were statistically significant using a Wilcoxon two-sample test with *t* approximation, which were then adjusted with a false discovery rate (FDR) calculation for multiple comparisons. **i) IFN-α**, HIV-positive young adults vs. HIV-negative young adults (p=0.0003), HIV-positive older adults vs. HIV-negative older adults (p=0.011). All other comparisons were not significant. **ii) TNF**-**α**, HIV-negative older adults vs. HIV-negative young adults (p=0.0085), All other comparisons were not significant. **iii) IL-12**, HIV-positive older adults vs. HIV-negative older adults (p=0.0011). All other comparisons were not significant. **iiii) IL-6**, HIV-positive older adults vs. HIV-negative older adults (p=0.035). All other comparisons were not significant.

To study the basis for altered Dectin-1-dependent cytokine production, we evaluated the effects of age and HIV infection on Dectin-1 surface expression in monocyte subsets (Fig 2). We found significantly increased surface expression of Dectin-1 in both older adults and HIV-infected individuals in all monocyte populations with the exception of the nonclassical (CD14+ CD16H) where we only saw an HIV-associated difference in Dectin-1 surface expression. For HIV-positive young adults, Dectin-1 expression was increased in all monocyte subsets compared to HIV-negative young individuals, and we found a similar increase in HIV-positive older, compared to HIV-negative older adults that was statistically significant for the CD14+ CD11b+ and CD14+ CD16H subsets. For the activated CD14+ CD11b+ monocytes, in which we saw both an age and HIV-associated increase in Dectin-1 surface expression, we built a multivariable regression model to incorporate the effects of clinical covariates (Table 2). In the unadjusted model, both age and HIV-infection were significant and non-interacting contributors. However, the addition to the model the co-variates of recreational drug use, history of fungal disease, number of medical co-morbidities, and percent lifespan with HIV led to a loss of significance for age. In the fully adjusted model, Dectin-1 surface expression was significantly associated with HIV-infection, percent lifespan with HIV, recreational drug use, and number of co-morbidities.

**Table 2.**
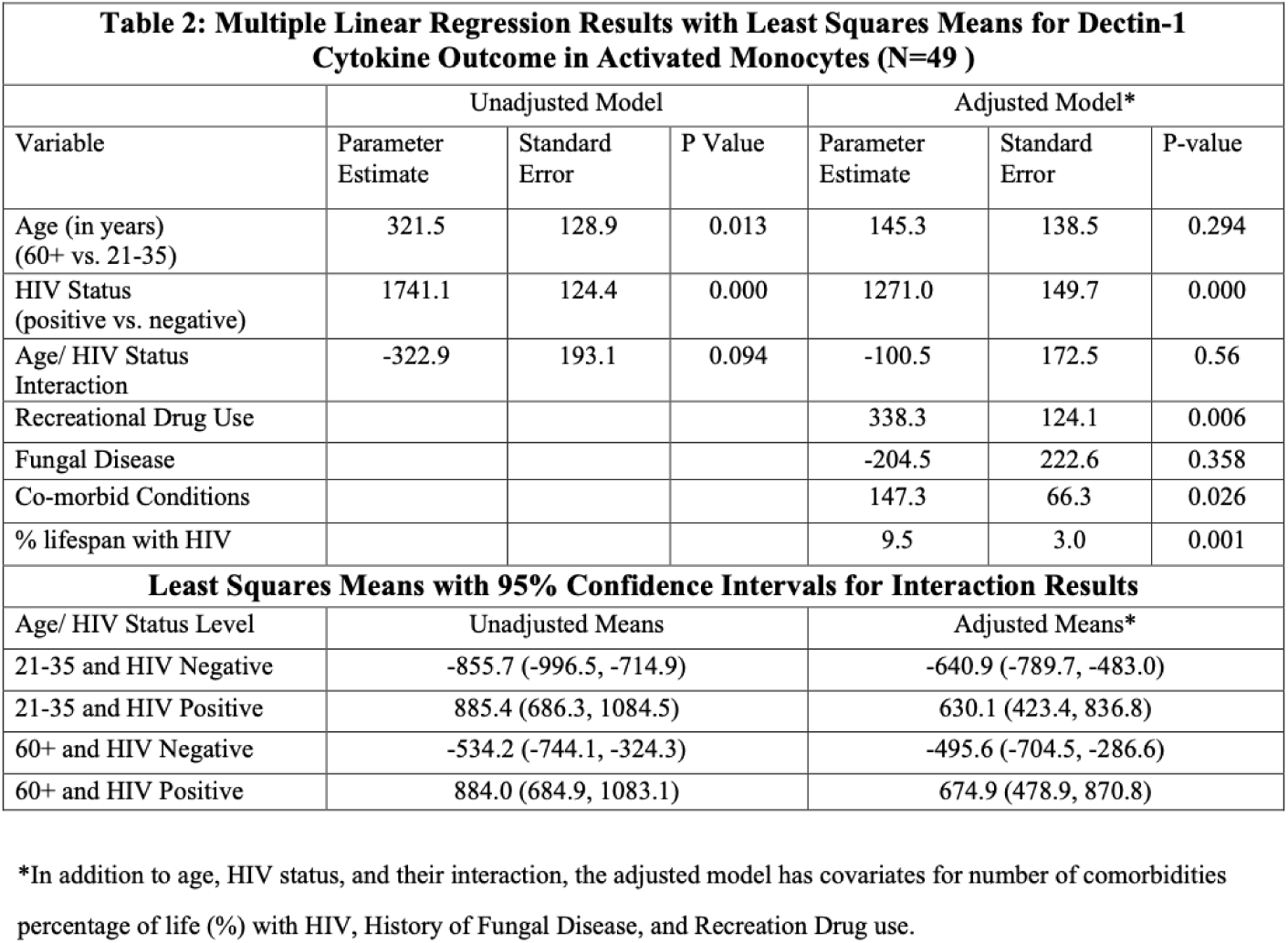
Dectin-1 surface expression in Activated Monocytes is significantly associated with HIV-infection, Recreational drug use, Co-morbid conditions, and % lifespan with HIV. Table outlining the multiple variable linear regression model for Dectin-1 surface expression, with the following variables (Age, HIV status, Age/HIV interaction term), and the following covariates (history of recreational drug use, history of fungal infection, number of co-morbid conditions, and % HIV life span).

**Figure 2.**
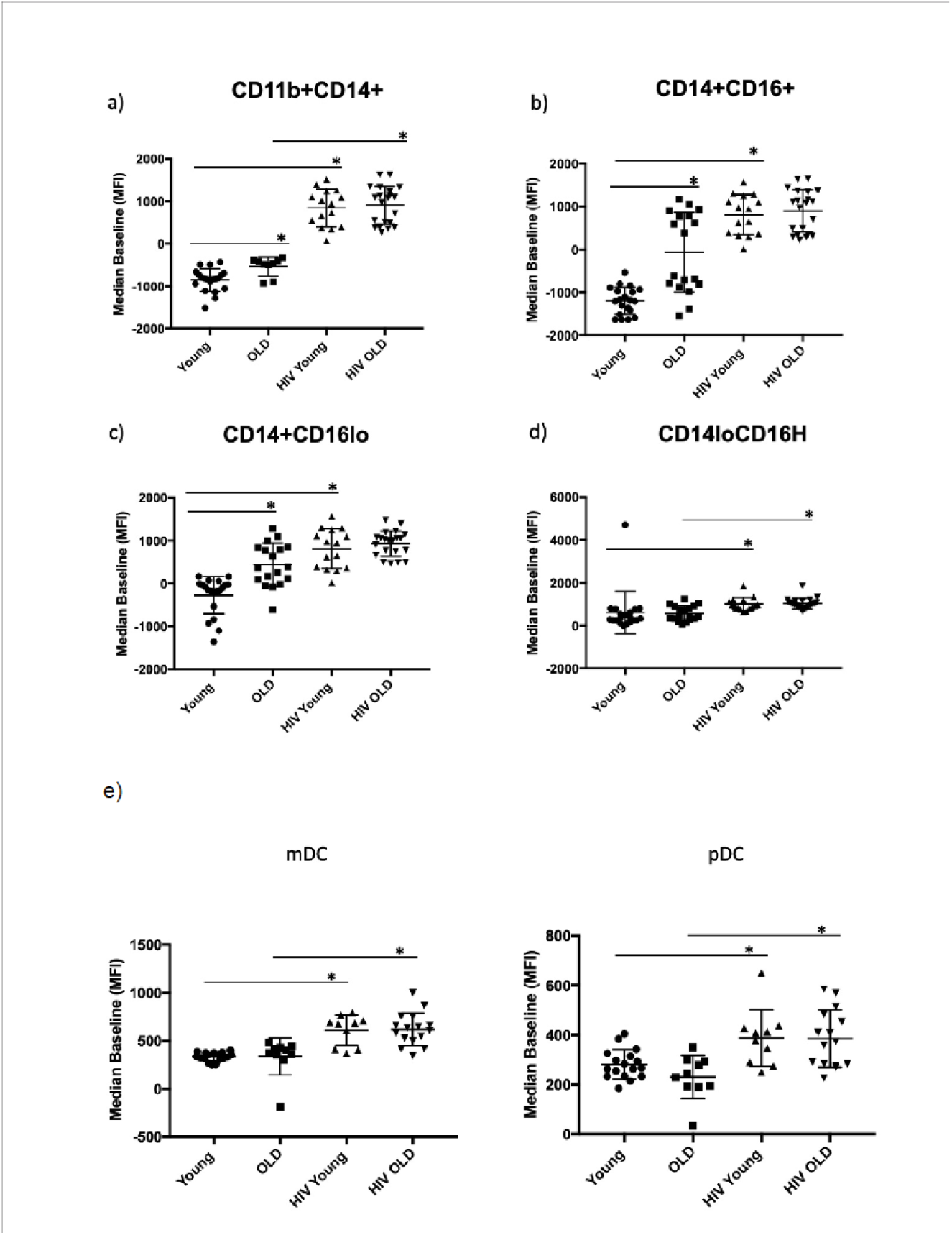
Dectin-1 Surface Expression in Monocytes and Dendritic cells. The cohort consists of the following: Monocytes: [HIV-negative adults (n=21), HIV-negative older adults (n= 24), HIV-positive young adults (n =14), and HIV-positive older adults (n = 22)]. Dendritic cells: [HIV-negative young adults (n=21), HIV-negative older adults (n=18), HIV-positive young adults (n=10), HIV-positive older adults (n= 17)]. Dot plots showing the Median of the Mean Fluorescence Intensity (MFI) of Dectin-1 surface expression. The following comparisons indicated by asterisks were statistically significant using a Wilcoxon two-sample test with *t* approximation, which were then adjusted with a false discover rate (FDR) calculation for multiple comparisons. **a) Activated Monocytes (CD11b+ CD14+)**. HIV-negative older adults versus young adults (p= 0.026), HIV-positive young adults versus HIV-negative young adults (p=0.001), HIV-positive older adults versus HIV-negative older adults, (p=0.005). All other comparisons were not significant. **b) Inflammatory Monocytes (CD14+CD16+)**. HIV-negative older adults versus young adults (p= 0.003), HIV-positive young adults versus HIV-negative young adults (p=0.001). All other comparisons were not significant. **c) Classical Monocytes (CD14+CD16lo)**. HIV-negative older adults versus young adults (p= 0.0039), HIV-positive young adults versus HIV-negative young adults (p=0.0026). All other comparisons were not significant. **d) Non-classical monocytes (CD14lo, CD16H)**. HIV-positive young adults versus HIV-negative young adults (p=0.011), HIV-positive older adults versus HIV-negative older adults, (p=0.048). All other comparisons were not significant. **e). Dectin-1 Surface Expression in Myeloid Dendritic cells (mDC) and Plasmacytoid Dendritic Cells (pDC)**. Dot plots showing the Median of the Mean Fluorescence Intensity (MFI) of Dectin-1 surface expression. The following comparisons indicated by asterisks were statistically significant using a Wilcoxon two-sample test with *t* approximation, which were then adjusted with a false discover rate (FDR) calculation for multiple comparisons. **i). mDC**, HIV-positive young adults vs. HIV-negative young adults (p=0.0002), HIV-positive older adults vs. HIV-negative older adults (p=0.0018). ii). **pDC**, HIV-positive young adults vs. HIV-negative young adults (p=0.012), HIV-positive older adults vs. HIV-negative older adults (p=0.0085).

We also examined the response of Dectin-1 stimulation in dendritic cells (DCs) in the context of aging and HIV-infection. Lineage negative and HLA-DR positive cells were separated into CD11c+, CD123-(mDCs), and CD11c-, CD123+ (pDCs) (Panda et al., 2010). We determined intracellular production of IFN-α, TNF-α, IL-6 and IL-12 in both subsets following WGP stimulation. mDCs (Figure 1c) showed a significant age-associated increase in IFN-α and TNF-α production with Dectin-1 stimulation, and an HIV-associated increase in IFN-α and IL-12 production in both young and older adults. IL-6 production was comparable among these groups except for a significant increase in HIV-positive versus HIV-negative older adults. Cytokine production in pDCs (Figure 1d) was comparable in HIV-negative young and older adults. However, there was a significant HIV-associated increase in IFN-α, TNF-α (in HIV-positive young) and IL-6 and IL-12 (in HIV-positive older) production. Dectin-1 surface expression levels in mDCs and pDCs showed a significant HIV-associated increase when compared to HIV-negative adults (Figure 2 b). Overall, our findings show that Dectin-1-dependent type I interferon production is increased with HIV infection in mDCs and pDCs from both young and older adults, with age-related increases in IFN-α and TNF-α found in mDCS from HIV-negative older adults.

### Effects of age and HIV infection on transcriptomic profiling identifies differentially expressed genes in inflammatory monocytes from the different cohorts

We investigated the basis for the age and HIV-associated alterations in Dectin-1 function, focusing on the CD14+ CD16+ inflammatory monocyte subset, which overlaps with the CD11b+ subset and is expanded in the setting of aging and HIV-infection (Zapata & Shaw, 2014a). Sorted CD14+ CD16+ monocytes from 8 HIV-positive and 11 HIV-negative young and older adults were stimulated with WGP and subjected to RNA-seq analysis. The clinical characteristics for all the enrolled individuals used for the RNAseq study can be reviewed in Supplemental Table 2.

Principal component analysis explained 69% of the variance; baseline and Dectin-1-stimulated inflammatory monocytes were clearly distinguished on PC2, while monocytes from HIV-positive and HIV-negative older adults were separated on PC1 (accounting for 53% of the variance). Notably, while samples from young HIV-positive vs. young HIV-negative adults did not show obvious separation (including HIV-negative young adults with increased comorbid medical conditions), there was a clear distinction between HIV-positive and HIV-negative older adults both at baseline and following Dectin-1 stimulation. Notably, HIV-older adults showed distinct separation from the other groups (Figure 3a).

**Figure 3.**
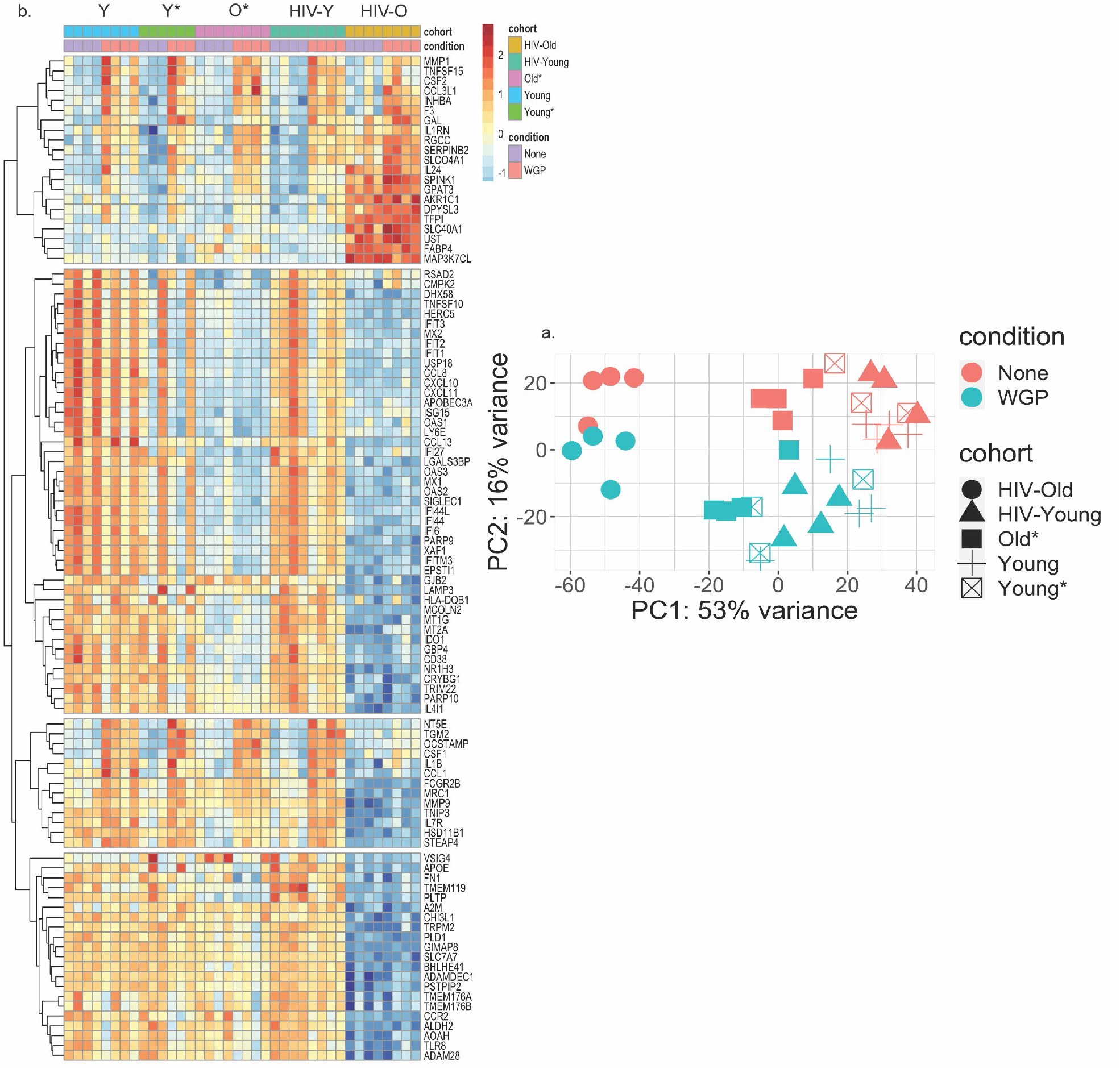
**a) Principal Component Analysis Highlights Transcriptional Differences between the HIV-positive Older Adults and All Other Cohorts**. Principal component analysis was performed to identify age, HIV, and Dectin-1 stimulation associated differences among Inflammatory (CD14+CD16+) monocytes. Dectin-1 stimulated vs. unstimulated Inflammatory monocytes were separated with 16% variance on PC2, and monocytes from HIV older adults vs all other cohorts are separated by 53%. [HIV-OLD n= 4, HIV-Young n =4, Young = 3, Young* = 4, Old = 3]. Please note that the asterisk (*) indicates the presence of co-morbidities. None = Unstimulated, WGP= Dectin-1 stimulation. b) **Heatmap of top 100 variable genes**. A total of 314 (Young (Y)), 541 (Young * (Y*)), 402 (Old (O)),860 (HIV-Young (HIV-Y)), and 156 (HIV Old (HIV-O)) unique DEG (differentially expressed genes) were noted for each of the cohorts respectively. The genes with a fold change of (FC) 1.2 and FDR ≤ 0.1 were defined as differentially expressed genes (DEGs). This heat map notes differences in HIV OLD vs all other cohorts– The effect of stimulation is seen in the second row (Pink column). The heatmap was constructed using Pheatmap. The transcripts were normalized using variance stabilizing transformation function. The color represents relative expression of transcripts that covary across cohorts and condition.

Upon WGP stimulation, we identified 314, 541, 402, 860, and 156 differentially expressed genes (DEGs) relative to baseline among Young, Young* (with increased medical comorbidities), Older, HIV-Young, and HIV-Older, respectively. A heat map of the top 100 variable genes among the cohorts is shown in Figure 3b, and shows baseline and post-stimulation differences in the HIV-negative young cohorts with and without increased medical comorbidities. Substantial alterations in gene expression were found for both HIV-negative and HIV-positive older, compared to young individuals.

### Dectin-1 Stimulation Elicits both Distinct and Common Transcriptional Gene Signatures in each cohort

Functional enrichment was performed on all the cohorts that were Dectin-1 stimulated. The Hallmark gene set enrichment analysis showed significant enrichment for the inflammatory response, and TNF-α and IL-2/STAT5 signaling in all cohorts in response to Dectin-1 stimulation with increased enrichment noted in the HIV older group and the Older HIV-negative group compared to the young adults (Figure 4). The functional analysis of DEGs using KEGG pathways (Supplemental Figure 3) confirmed significant enrichment of TNF signaling in all cohorts, in agreement with Hallmark gene sets. KEGG analysis also showed upregulation of IL-17 signaling, cytokine-cytokine receptor interactions, NF-κB signaling, complement & coagulation cascades, and MAPK signaling pathways in response to Dectin-1 stimulation in most cohorts (Supplemental Figure 3).

**Figure 4.**
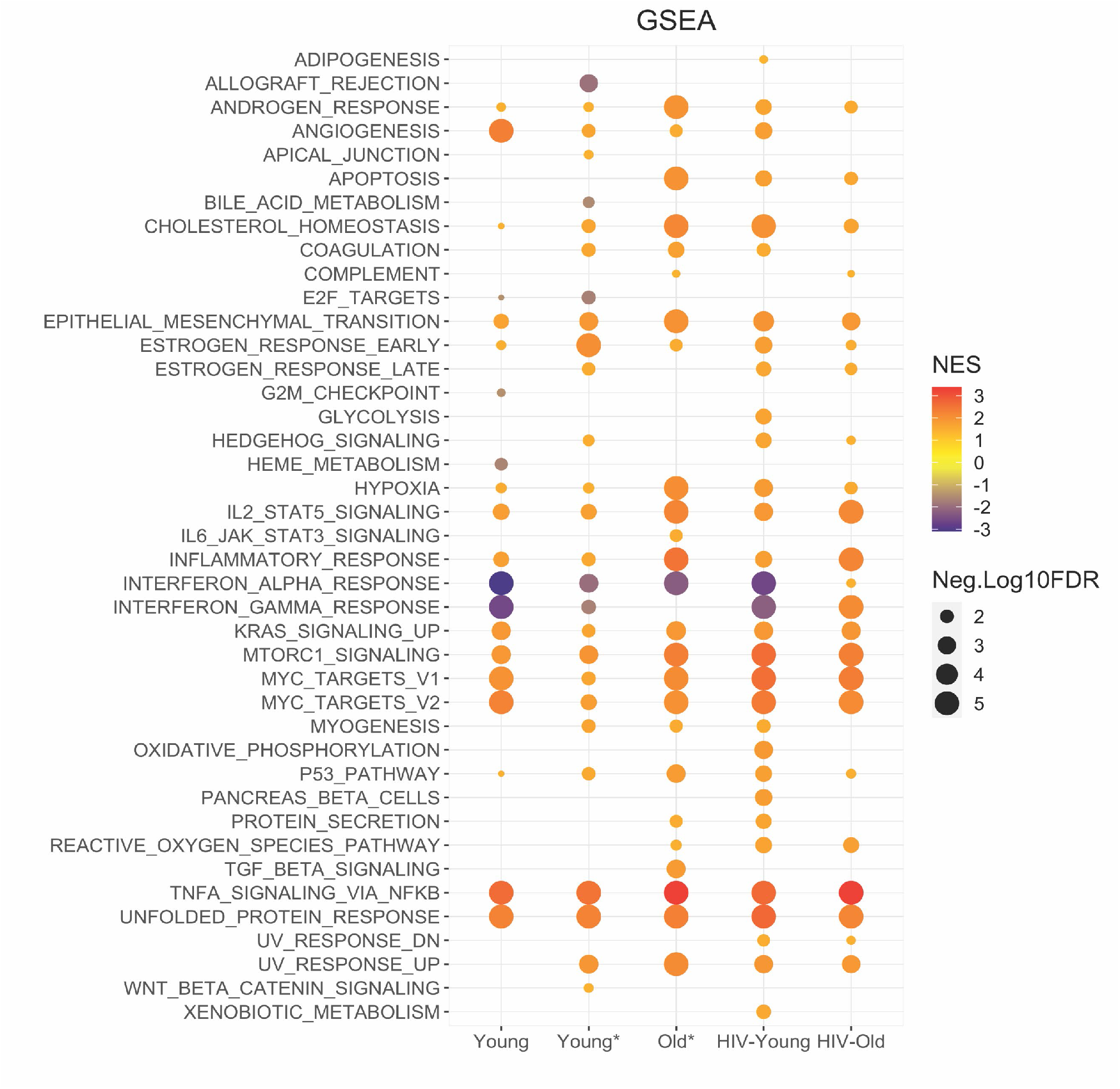
Enrichment Analysis Highlights Unique Signatures with Dectin-1 Stimulation Across Cohorts. Functional enrichment was performed using the Broad institutes Hallmark gene set enrichment analysis. GSEA performed using Hallmark gene sets for each cohort. The dot graph represents the significant Hallmark pathways identified in WGP stimulated Inflammatory monocytes of the respective cohort. The pathways with FDR of <u><5</u>% were considered significant. For graphical representation, FDR values with 0 were adjusted to 0.00001. The size of the node represents -Log10(FDR) and color of node represent normalized enrichment score. NES = Normalized enrichment scores.

Several metabolic pathways were also strongly induced with Dectin-1 stimulation, including mTORC1 signaling and the hypoxia pathway in all cohorts (Fig 4) with increased enrichment of mTORC1 signaling noted in the setting of aging and HIV-infection. With respect to the PI3K-Akt signaling pathway; we found diminished in PI3K-Akt signaling in the HIV Older group, that could in part reflect baseline pathway enrichment relative to young HIV-negative adults (Supplemental Figure 3 and Supplemental Figure 6). GSEA baseline analysis also showed upregulation of mTORC1 signaling in the HIV older group that was not seen in any other cohort (indeed downregulation was found in the HIV young group) (Supplemental Figure 7). Finally, the unfolded protein response, indicative of endoplasmic reticulum stress was strongly upregulated in all cohorts with Dectin-1 stimulation (Fig. 4). Overall, these findings suggest that age and HIV infection potentiate Dectin-1-dependent inflammatory signaling in CD14+ CD16+ monocytes and that HIV infection in older adults may result in dysregulated metabolic signaling via the mTORC1/PI3K pathway.

The HIV-older group in particular showed additional differences in gene expression. In contrast to young adults, both HIV-older and HIV-negative Older* adults failed to upregulate PPAR and Rap1 signaling pathways in response to Dectin-1 stimulation (Supplemental Figure 3). In this regard, KEGG Baseline pathway analysis showed that PPAR signaling was already upregulated in the HIV-Older adults (Supplemental Figure 8). Besides regulating cytokines, Dectin-1 stimulation is also known to initiate the production of reactive oxygen species (ROS) (Gantner, Simmons, Canavera, Akira, & Underhill, 2003). Dectin-1 stimulation of inflammatory monocytes resulted in the significant enrichment of the Hallmark Reactive Oxygen Species pathway among Older, HIV-Older and HIV-young but not young HIV-negative adults (Fig. 4), with baseline upregulation of the ROS pathway noted only in HIV-Older adults (Supplemental Figure 7). While cytokine signaling pathways were generally upregulated in stimulated inflammatory monocytes (Supplemental Figure 4a, d), HIV-Older individuals demonstrated higher baseline levels for the TNF signaling pathway, complement coagulation cascades (Supplemental Figure 4b, c) and IL-19 and IL-24 (Supplemental Figure 4a, d) gene expression compared to other cohorts. IL-7R was upregulated in all cohorts except for the HIV-Older cohort (where upregulation was minimal or absent). Similarly, IL1-β and IL-1A baseline expression was higher in all young cohorts compared to the HIV-positive and HIV-negative older adults, where baseline levels were absent (Supplemental Figure 4a, d). Induction of IL1-β and IL-1A gene expression with stimulation was seen in all cohorts, although less so in the HIV-Older cohort.

### Dectin-1 induces Interferon response in HIV-Older adults

GSEA analysis exhibited downregulation of the Interferon-α and -γ response in inflammatory monocytes from all cohorts except for the HIV Older cohort (Figure 4), in which upregulation was observed. Interestingly, this gene expression data parallels the increased production of IFN-α that was noted in the HIV Older cohort in both myeloid dendritic cells and plasmacytoid dendritic cells (Figure 1c, d). Further examination of these gene sets demonstrated increased basal levels of the IFN--α and -γ response in young adults (both HIV-negative and HIV-positive) that was downregulated (Supplemental Figure 5a) upon Dectin-1 stimulation; in contrast, both HIV-negative and HIV-positive older adults showed decreased basal IFN expression that increased with stimulation. In HIV-older adults (Supplemental Figure 5b) IFN-α and IFN -γ were predicted as an activated upstream regulators of the following IFN inducible genes including; CMPK2, RSAD2, RIPK2, GZMB, CD274 (PDL-1) further confirming downstream signaling.

### Activation of Dectin-1/CLEC-7 signaling in Inflammatory Monocytes Across Cohorts

As expected, Gene set enrichment analysis (GSEA) for REACTOME demonstrated significant enrichment of the reactome Dectin-1 mediated non-canonical NF-κB gene set and Dectin-1 (CLECA7A) signaling pathways gene set in Dectin-1 stimulated Inflammatory monocytes of HIV-Young, HIV-Older, and Older HIV-negative adults (Supplementary Figure 9 a). Enrichment scores and q-values were highly significant for the HIV-older and the older (with co-morbidities) (NES>2, q<0.001) HIV-negative group compared to that of HIV-Young adults (Supplemental Figure 9a). Enrichment of Dectin-1 signaling was not significant in the young HIV-negative cohorts (Young and Young*). Although, upregulation of Dectin-1 signaling genes, BCL10 and MALT1 was observed in all cohorts following WGP stimulation of monocytes (Supplemental Figure 9b). Bcl10 and MALT1 both form part of the CARD9 complex that is important for downstream Dectin-1 signaling (Plato et al., 2013).

### TREM-1 enrichment is associated with Dectin-1 activation

We also carried out GSEA using ImmuneSigDB gene sets, and observed significant enrichment (NES>3, q<0.0001) in Dectin-1 stimulated monocyte subsets (Supplemental Figure 9a) in all groups for Triggering receptor in myeloid cells (TREM-1), an innate immune receptor with unknown activating ligand that has been shown to modulate the inflammatory response (Bouchon, Facchetti, Weigand, & Colonna, 2001; Feng et al., 2021; Gallop et al., 2021; Zhong et al., 2016). Our data also demonstrates that TREM-1 is a prominent upstream regulator of the response to β-D-glucans (Figure 5b) as noted in the next section.

**Figure 5.**
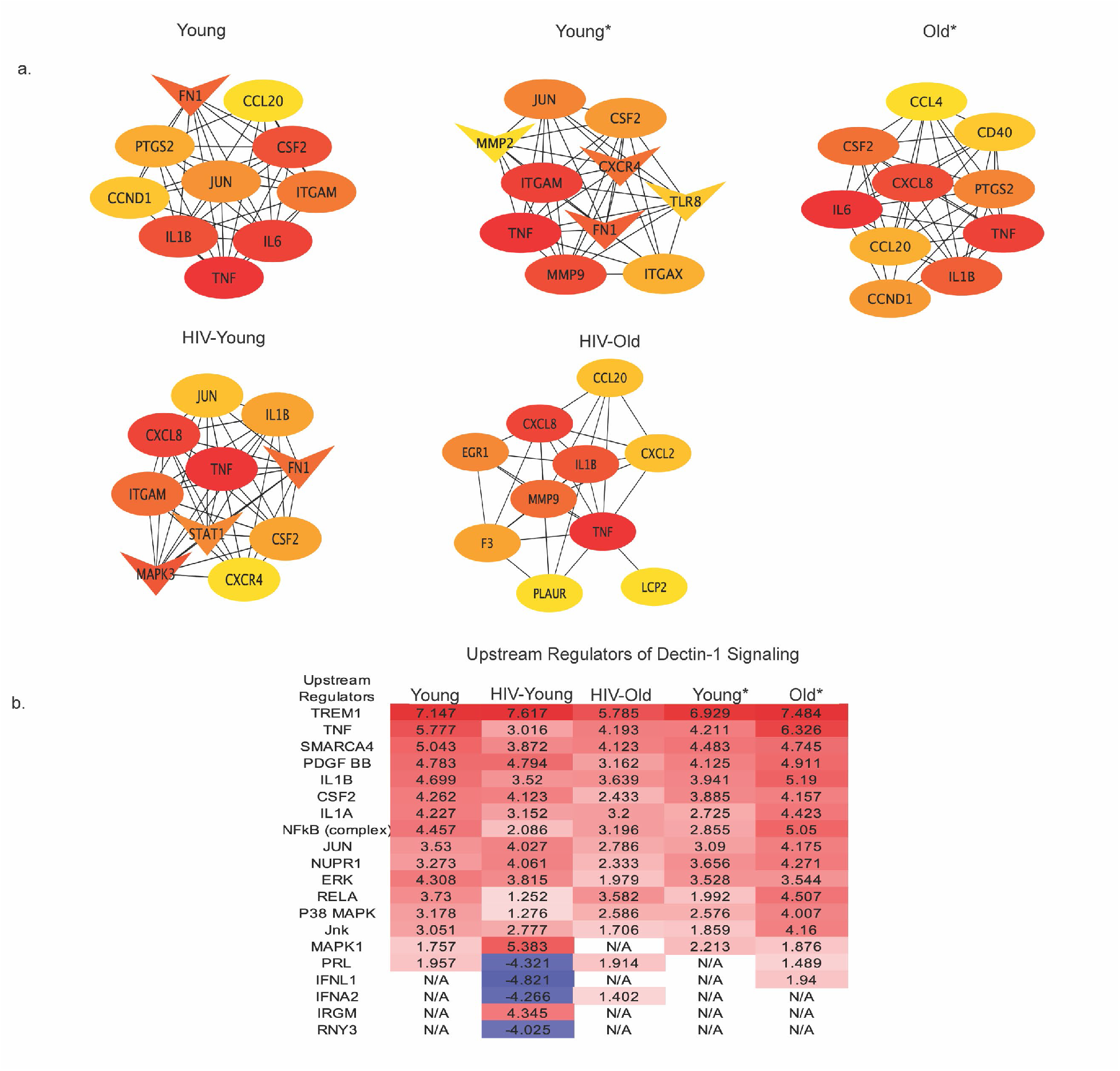
Upstream Regulators and Hub Genes in Dectin-1 signaling. **a. Hub Gene Networks**. Protein-protein interaction network of hub genes identified based on the degree by using CytoHubba plugin of the Cytoscape. Hub genes were identified in all cohorts, Young, Young*, Old*, HIV-Young and HIV-Old. The red color of the node represents a higher degree of interaction, orange represents an intermediate degree of interaction, and nodes with yellow color represent a lower degree of interaction compared to others. Oval nodes reflect upregulated DEGs, triangle shaped nodes represent down-regulated DEGs. **b. Upstream Regulators of Dectin-1 Signaling**. DEGs were analyzed for upstream regulators using IPA and top upstream regulators (including transcription factors) with Z-scores cutoff of 3 are listed for all cohorts.

### Hub Genes and Upstream Regulators

The DEGs were further analyzed for protein-protein interaction (PPI) networks by using STRING, and Cytoscape. [The Hub genes were identified from PPI network based on their degree using Cytohubba plugin of Cytoscape.] The top ten hub genes identified in inflammatory monocytes in response to Dectin-1 stimulation are shown in Figure 5a. The common hub genes in many of the cohorts included TNF-α, IL1-β, CSF2 (colony stimulating factor 2), CXCL8 (IL-8), CCL20 (MIP3-α), JUN and ITGAM (CD11b). Hub genes that were unique to the HIV-Older cohort included the coagulation factors F3 (tissue factor) and PLAUR (plasminogen activator), EGR1 (Early growth response protein 1), and LCP2 (Lymphocyte Cytosolic protein 2). Interestingly, the downregulation of STAT1 was only observed among HIV-Young individual cohort. These differences in Hub genes imply differential unique protein interactions in each of the cohorts with respect to Dectin-1 signaling.

We also analyzed DEGs for the prediction of upstream regulators by using ingenuity pathway analysis (IPA). The top upstream regulators with Z-scores cutoff of 3 are listed in Figure 5b, while the complete list of the significantly predicted upstream regulators are provided in Supplementary Table 1. The top most activated upstream factors in response to Dectin-1 stimulation includes TREM1, TNF, SMARCA4 (actin dependent regulator of chromatin, subfamily A, member 4), IL-1β, NFκB, JUN, ERK, RELA and p38MAPK. The downregulation of the hub gene STAT1 (Figure 5b) along with the inhibition of upstream regulators, IFNL1 and IFNA2 (Figure 5b), suggests the Dectin-1 stimulation inhibits the interferon response in HIV-Young individuals. Both analyses (hub gene analysis and upstream regulators) suggest that TNF-α and IL1-β are central players of Dectin-1 signaling. Upstream regulator analysis shows MAPK signaling in general (p38, JUN, JNK, ERK, MAPK1) plays an important role in Dectin-1 stimulation across cohorts.

### Dectin-1 Stimulation of Monocytes Induces Distinct Differentiation Signatures

We observed significant enrichment of DEGs for transcriptional dysregulation in cancer and hematopoietic cell lineage KEGG gene sets along with significant enrichment for Hallmark epithelial mesenchymal transition (GSEA) in response to Dectin-1 stimulation (Figures 4 and Supplemental Figure 3). The activation of genes such as NR4A3, MET, ZEB1, CSF1, CSF2, CSF3, which can be involved in transcriptional dysregulation in cancer or hematopoietic cell lineage signaling pathways, can also be involved in cell differentiation (Boulet et al., 2019; Q. Chen, DeFrances, & Zarnegar, 1996; Scott & Omilusik, 2019). Therefore, we further investigated GO terms that could be associated with monocyte differentiation.

During infection, monocytes can terminally differentiate either to monocyte derived macrophages (moMac) and/or to monocyte-derived dendritic cells (moDCs). Recently, Boulet et al. 2019 demonstrated the importance of the orphan nuclear receptor NR4A3 in the differentiation of murine moDCs in response to LPS stimulation (Boulet et al., 2019). In our study, Dectin-1 stimulation of Inflammatory (CD14+CD16+) monocytes resulted in the significant upregulation of the NR4A3 and NR4A1 genes in all cohorts (Figure 6a). The expression of granulocyte-macrophage colony-stimulating factor (GM-CSF or CSF2) and macrophage-CSF (M-CSF or CSF1), required for macrophage/dendritic cell differentiation, were also significantly upregulated in all cohorts except for HIV-Old, where a trend was noted.

**Fig 6.**
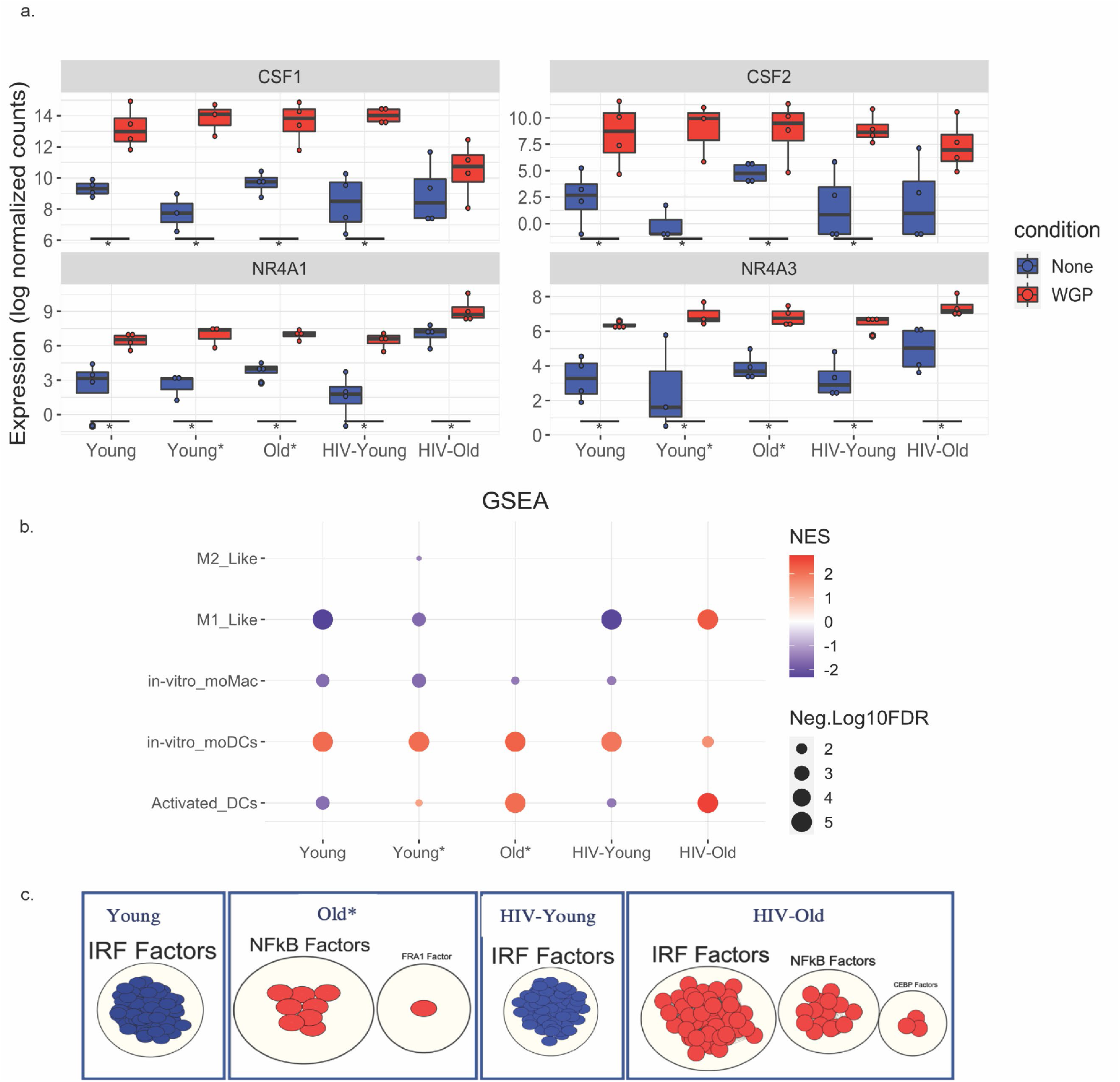
Dectin-1 stimulation leads to distinct signatures of differentiation. GSEA was performed on normalized counts obtained from Dectin-1 stimulated Inflammatory monocytes using previously defined gene sets for in vitro moDCs and Activated DCs. a) Box plots of CSF1, CSF2, NR4A1, and NR4A3. The symbol * represents significant differentially expressed genes with fold cutoff of 1.2, and q value < 0.1. b) Gene enrichment analysis for monocyte differentiation was performed. The plot represents normalized enrichment scores and FDR(q-values) obtained for each cohort. For graphical representation, FDR values with 0 were adjusted to 0.00001 and FDR of <5% was considered significant for the enrichment of a particular term. c) Transcription factor analysis of the significantly enriched activated DCs gene sets using gProfiler and visualization by Cytoscape. * Significantly differentially expressed genes with FC cut off 1.2 and padj <0.1.

Therefore, we explored if Dectin-1 stimulation regulates the differentiation or activation of either macrophages or dendritic cells. Gene set enrichment analysis using previously defined gene sets for monocyte differentiation (Banchereau et al., 2014; Goudot et al., 2017; Tang-Huau et al., 2018) (Becker et al., 2015), revealed significant enrichment for the *in vitro* monocyte derived dendritic cell gene set in Dectin-1 stimulated inflammatory monocytes of all cohorts (Figure 6b). The *in vitro* monocyte derived macrophage gene set was downregulated in most cohorts. However, Dectin-1 stimulated inflammatory monocytes from HIV-older adults showed significant positive enrichment for activated DC and M1-like gene sets. The activated DC gene set was also found to be significantly enriched in monocytes of young and older adults with co-morbid conditions (and downregulated in young HIV-negative and young HIV-positive adults. The gene sets for activated DCs were further analyzed with transcription factor (TF) analysis using gProfiler (Figure 6c), which demonstrated that the genes for activated DCs were positively regulated by Interferon Regulatory Factors (IRFs) in the HIV Older group, and negatively regulated by IRFs in both the HIV-negative and HIV-positive young adults. NF-κB transcription factors also showed positive regulation in the HIV Older group and in the Older HIV negative group.

Dectin-1 induced transcriptomic changes in inflammatory monocytes suggest the differentiation of CD14+CD16+ monocytes mainly to monocyte derived dendritic cells, as these changes were noted in all the cohorts. However, the transcriptomic profile of the stimulated inflammatory monocytes of HIV-older adults also revealed a distinct gene signature that demonstrated an M1-like phenotype in macrophages, as well as an activated DC signature that was also noted in young and Older HIV-negative adults with co-morbidities. Young HIV-negative adults seemed to skew towards a monocyte-derived dendritic cell signature. Overall, these results demonstrate that unique inflammatory signatures are associated with age, and HIV-infection status.

## Discussion

To our knowledge, our study is the first to evaluate the specific function of Dectin-1 in the setting of aging and HIV-infection. A few themes emerged from our studies. Stimulation of both monocytes and dendritic cells with β-D-glucans led to a more pro-inflammatory phenotype in both monocytes and dendritic cells in HIV-infected individuals that included increased levels of IL-12, TNF-α, and IFN-α. While this pro-inflammatory phenotype was evident in certain cell types (CD11b+ activated monocytes, myeloid dendritic cells) for HIV-negative older adults, the most striking differences were associated with HIV-infection. Our previous studies of Mincle, another C-type lectin receptor family member recognizing carbohydrate motifs including M. tuberculosis cord factor, also showed an increase in cytokine production in monocytes with age and HIV infection (Zapata et al., 2019); thus HIV and age appear to enhance cytokine production downstream of C-type lectin signaling and represent an additional factor that could contribute to age-related chronic inflammation.

Our analyses of RNA-seq experiments on purified CD14+ CD16+ monocytes from young and older adults with and without HIV infection were notable for a distinct inflammatory signature in the HIV-older population. Gene expression from several pro-inflammatory pathways was enhanced at baseline in this group, such as TNF-α signaling (and in this regard, Dectin-1-dependent TNF-α production in inflammatory monocytes was increased with HIV infection in both young and older adults (Fig. 2)). Increased GSEA enrichment of the IL-2/STAT5 signaling pathways was also noted in the HIV-positive Older cohorts (also seen in HIV-negative Older cohort). Additionally, there was also increased baseline expression of IL-19/IL-24 (Supplemental Figure 4 a), and the KEGG complement/coagulation pathways (Supplemental Figure 4c). The link between Dectin-1 and coagulation factors is unclear, but both F3 (tissue factor and PLAUR (plasminogen activator) were hub genes only for the HIV-Older group (Fig. 5a). Because increased production of clotting factors is also associated with age-associated chronic inflammation (Zapata & Shaw, 2014a), our results suggest CLR signaling as an additional contributing factor. Finally, increased Reactive Oxygen species (ROS) pathway signaling was noted in the Older HIV-negative and HIV-positive cohorts (with stimulation) (Figure 4). However, baseline analysis only showed increased enrichment of the ROS signaling in the HIV-Older cohort (Supplemental Figure 7). Overall, HIV-infected older adults demonstrated unique gene expression with Dectin-1 stimulation when compared to the other cohorts.

Studies in the early years of the HIV pandemic reported higher plasma levels of IFN-α in HIV-infected individuals in both acute and chronic infection (Bosinger & Utay, 2015) that was strongly associated with the immune activation seen with chronic HIV-infection and is thought to contribute to pathogenesis. Our studies indicate that Dectin-1 signaling, particularly in HIV-infected individuals, may contribute to this dysregulated IFN production. Notably, both mDCs and pDCs showed an HIV-associated increase in Dectin-1-induced IFN-α production in young and older adults (Fig. 1c and 1d). This increase in IFN-α production also paralleled the increased Dectin-1 surface expression in both mDCs and pDCs noted with HIV-infection (Fig. 2b). We also found increased enrichment of Dectin-1-dependent IFN-α and IFN-γ pathway expression only in the HIV-Older group, in contrast with downregulated IFNα and IFN-γ responses in all other cohorts (Fig. 4 and Supplemental Fig 5). In this regard, we found an age-associated decrease in basal IFN-α and IFN-γ pathway expression that may contribute to enhanced Dectin-1-dependent IFN signaling in HIV-Older adults. While traditionally associated with antiviral immunity, interferon responses have been implicated in antifungal immune responses (del Fresno et al., 2013; Inglis, Berkes, Hocking Murray, & Sil, 2010) (Smeekens et al., 2013). However, type I IFN production may also result in dysregulated inflammatory responses in monocytes that lead to worsened outcomes from fungal infection (Majer et al., 2012) as can be seen in older adults with HIV infection. Additionally, the increased and persistent production of IFNs via Dectin-1 stimulation may also be contributing to the pro-inflammatory environment that is seen in HIV-infection and Aging.

Analysis of our RNA-seq data revealed the innate immune receptor Triggering receptor in myeloid cells-1 (TREM-1) as a significant upstream regulator in all cohorts (Fig 5b). There are limited studies demonstrating a relationship between TREM-1 and Dectin-1 in fungal pathogenesis, although expression of TREM-1 and Dectin-1 both increased with severity and progression of an *Aspergillus* keratitis mouse model (Zhong et al., 2016), with inhibition of both TREM-1 and Dectin-1 resulting in protection from fungal disease. TREM-1 is a receptor expressed on neutrophils and monocytes that modulates the inflammatory response (Arts, Joosten, van der Meer, & Netea, 2013; Tammaro et al., 2017), and is upregulated in sepsis. Although, the specific agonist for TREM-1 remains unknown, the receptor is known to interact with CARD9 in a complex with Bcl-10 and MALT-1, hinting at a possible interaction with C-type lectin receptors (Arts et al., 2013). The significant upregulation of TREM-1 noted in our study also alludes to a physical interaction between TREM-1 and Dectin-1.

Other upstream regulators that emerged included the cytokines TNF-α, IL-1β that were significant in all cohorts, and IFNs that were prominent negative regulators in the HIV-Young cohort. Other important regulators that emerged was the NF-κB complex, and the MAP kinases ERK, and p38 pathways which have been reported as important components of the Dectin-1 signaling (Elcombe et al., 2013; Plato et al., 2013). In fact, ERK and p38 were shown to regulate zymosan induced IL-10 production in a mouse model (Elcombe et al., 2013). IL-10 was robustly produced by all cohorts in our study in response to Dectin-1 stimulation. Dectin-1 stimulation also increased expression of the anti-inflammatory cytokines IL-19 and IL-24 in the CD14+CD16+ monocytes in most cohorts, with the HIV-Older cohort already increased at baseline (Supplemental Figure 4a).

Other notable differences should be pointed out. The HIV-Older group showed decreased gene expression of IL-7R gene expression with Dectin-1 stimulation when compared to all other cohorts (Supplemental Figure 4a). Expression of IL-7R in monocytes has been reported in the setting of LPS exposure and its role remains to be fully defined (Al-Mossawi et al., 2019). Decreased expression of IL-7R in monocytes has not been reported in the setting of HIV and aging. However, decreases in IL-7R expression and impaired IL-7 signaling in T cells has been reported in the setting of both aging (Ucar et al., 2017) and HIV-infection (Bazdar, Kalinowska, & Sieg, 2009). Another gene of note was IL-1β which was noted to be increased at baseline in all cohorts except for the HIV-Older cohort (Supplemental Figure 4a). Dectin-1 stimulation led to an upregulation of IL1-β, in all cohorts (Supplemental Figure 4a,d), although higher in all groups compared to the HIV Older group. Dectin-1 is a known inducer of the inflammasome (Gringhuis et al., 2012; Kankkunen et al., 2010). Interestingly, IL-1β emerged as a significant upstream regulator of Dectin-1 signaling in all cohorts, in addition to an important hub gene (most cohorts) as noted in Fig 5b and 5a. Further study of the Dectin-1 induced inflammasome is needed in the setting of HIV-infection and aging.

Overall, Dectin-1 stimulation led to the upregulation of several metabolic pathways, including mTORC1 signaling, the hypoxia pathway, and P13K-Akt signaling. While upregulation of the MTORC1 signaling and the Hypoxia pathway was noted all cohorts (Fig 4), increased MTORC1 signaling was noted with both aging and HIV-infection. Baseline GSEA pathway analysis also demonstrated increased MTORC1 pathway enrichment in HIV-Older adults which was absent in all other cohorts. (Supplemental Figure 7). KEGG gene enrichment analysis showed upregulation of PI3K-Akt pathway in all groups except the HIV Old cohort (Supplemental Fig 3), although was also noted to be upregulated at baseline (Supplemental Figure 6). Activation of the Akt/mTOR/HIFβ−α pathway is thought to be the metabolic basis for trained immunity in Monocytes trained with β−glucan (Cheng et al., 2014), and trained immunity is likely playing a role in the anti-fungal response in humans. Our findings show that trained Immunity may also be contributing to the sustained pro-inflammatory phenotype seen in the setting of HIV and aging, a finding which has also been noted in other studies (van der Heijden et al., 2021) Furthermore, Dectin-1 stimulation also led to a strong upregulation of the Unfolded protein response in all cohorts (Fig 4). There is evidence that the unfolded protein response may be important for proper innate immune function and may be necessary for the proper functioning of dendritic cells (Todd, Lee, & Glimcher, 2008).

Finally, Dectin-1 led to the upregulation of several differentiation signatures (Fig 6) alluding to the possibility that Dectin-1 stimulation may lead to the differentiation of monocytes into both dendritic cells and macrophages. Specifically, Dectin-1 induced transcriptomic changes in monocytes suggest the differentiation of CD14+CD16+ monocytes mainly to monocyte-derived dendritic cells--changes noted in all the cohorts (Fig 6b). However, the transcriptomic profile of the stimulated inflammatory monocytes of HIV-positive older adults also revealed a distinct gene signature that demonstrated an activated DC signature, as well as a more inflammatory M1 macrophage signature. A prominent activated DC signature was also seen in Older HIV-negative adults with co-morbidities (a smaller upregulation was noted in Young HIV-negative adults with co-morbidities). Young HIV-negative adults seemed to skew towards a non-activated dendritic cell signature. Activation of the PPAR-γ pathway has been shown to inhibit Dectin-1 induced dendritic cell activation in human PBMCs (Kock et al., 2011) from healthy volunteers. Interestingly, KEGG DEG analysis did not show upregulation of the PPAR signaling pathway in the HIV-Older and Older HIV-negative group as compared to young adults where it was upregulated. However, baseline pathway analysis also showed upregulation of PPAR signaling at baseline in the HIV Older cohort—alluding to more complex signaling in the setting of HIV and aging. Overall, Dectin-1 signaling in the setting of HIV and Aging was associated with a more inflammatory activated DC signature and M1 macrophage signature.

Other roles for Dectin-1 have emerged which may also be playing a role in the setting of HIV-infection and aging. There is accumulating evidence that Dectin-1 recognizes endogenous ligands such as annexins, and vimentins from dying cells (Deerhake & Shinohara, 2021). Vimentin when bound to Dectin-1 in atherosclerotic plaques induces ROS production in human monocytes (Thiagarajan et al., 2013). Dectin-1 activation induced by vimentin promoted obesity and insulin resistance in a diet-induced obesity mouse model. This same study showed increased Dectin-1 expression in human adipose tissue samples of obese individuals when compared to lean subjects (Castoldi et al., 2017). Interestingly, a recent study of caloric restriction in humans showed a downregulation of Dectin-1 signaling in RNA seq studies of subcutaneous adipose tissue after two years of 14% caloric restriction (Spadaro et al., 2022). In our study, Dectin-1 surface expression in activated monocytes was examined in the context of a multivariable regression model (Table 2). Within our fully adjusted model, the co-variates of HIV-infection, recreational drug use, number of co-morbidities, and percent lifespan with HIV all proved to be significant. The significance of number of co-morbidities in the model of Dectin-1 surface expression suggests that factors unrelated to fungi are important for Dectin-1 surface expression and need to be further explored. Therefore, the stimulation of Dectin-1 by endogenous ligands could certainly be contributing to both age and HIV-associated chronic inflammation as seen in our study.

In addition to endogenous ligands, circulating fungal ligands could also be contributing to these notable age and HIV-associated differences. Elevated levels of circulating β-D-glucan levels have been noted in several studies in the setting of chronic HIV-infection that persist despite long term Antiretroviral therapy (ART) and is thought to be secondary to microbial translocation (Hoenigl et al., 2016; Mehraj, Ramendra, & Isnard, 2020). Previous studies have shown that elevated levels of β-D-glucan was associated with markers of immune activation (soluble CD14, HLA-DR+ CD4/CD8 T cells, and IL-6/IL-8 levels) and decreased Dectin-1 surface expression on monocytes from individuals with HIV infection, compared to HIV-negative controls (Mehraj et al., 2020). HIV-positive and negative individuals in this study were not segregated into age groups, and so it is not clear if the general HIV-associated increase in Dectin-1 expression we observed would have been found; in addition, there were methodological differences between our studies, such as the analysis of freshly isolated PBMCs (this study), versus cryopreserved samples. Our findings showed increased Dectin-1 surface protein expression with both increased age and HIV-infection. Evaluation of Dectin-1 (Clec7a expression) gene expression (Supplemental Figure 10) showed an opposite trend of decreased gene expression levels in the HIV-Older cohort and increased levels in the young HIV-negative cohort. These findings suggest a mechanism that is regulated post-transcriptionally in the HIV-Older cohort that needs to be further investigated. A recent paper that examined monocytes from people living with HIV found a strong correlation between serum levels of β-D-glucans and production of IL-1β in general (although specific stimulation with Candida albicans led to decreased cytokine production in this study) and suggested that the robust production of cytokines by monocytes in HIV-infection was linked to trained immunity in monocytes (van der Heijden et al., 2021). Microbial translocation is something that is also likely occurring with increased age (Stehle et al., 2012) as increased levels of surrogate markers of microbial translocation have been noted in older adults. Therefore, our findings of a unique pro-inflammatory signature in the setting of HIV and aging could be secondary to elevated levels of circulating of β-D-glucans that have been noted in the setting of chronic HIV-infection (Zapata & Shaw, 2014a).

Overall, this study demonstrates that age, HIV-infection and co-morbidities can alter the individual immune response. In particular our study showed a unique immune signature in the setting of both HIV and Aging in response to Dectin-1 stimulation. As our HIV-positive population ages these immune differences will need to be considered for future vaccines and therapeutics.

## Methods

### Clinical Study Design and Recruitment of Participants

We recruited a total of 81 HIV-negative and HIV-positive participants, divided between young (21-35) and older (≥ 60 years) adults (Note: the age range for the Dendritic cell experiments was extended to 21-40 and ≥ 50 years). Subjects were recruited from the Yale Health Services, Yale Primary Care Center (PCC), and the Nathan Smith HIV Clinic. Participants were evaluated for clinical characteristics by chart review and self-report for demographic information, medications, CD4 count, HIV viral load, and comorbid conditions. Participants were excluded for the following reasons: an acute infection or antibiotic use within two weeks of recruitment, pregnancy, history of current cancer, history of stem cell, bone marrow or solid organ transplant, cirrhosis of the liver, kidney disease requiring dialysis, immunodeficiency other than HIV, and active Hepatitis B or C infection.

### Sample Preparation

Peripheral blood mononuclear cells (PBMCs) were isolated from heparinized blood using Ficoll gradient centrifugation (Histopaque Sigma) as previously described (Panda et al., 2010; van Duin et al., 2007). Freshly isolated PBMCs were stimulated with 10 μg/ml of whole glucan particles (WGP Dispersable or 1,3/1,6-β-glucan) (InvivoGen, tlr-wgp) which is a specific Dectin-1 agonist that lacks TLR stimulating activity--for 18 hours (for Monocyte experiments), and 12 hours (for dendritic cell experiments) in RPMI medium supplemented with 10% fetal bovine serum and Penicillin/Streptomycin/L-Glutamine at 37 °C. WGP (Soluble, or 1,3/1,6-β-glucan) (Invivogen tlr-wgps) was used as a negative control. WGP soluble particles can bind to the Dectin-1 receptor without activating it. Brefeldin-A (GolgiPlug BD Biosciences) was added for the last 6 hours of stimulation. Cells were then surface stained with anti-CD14-PE-CF594 (3G8, BD-Biosciences), anti-CD16-PE-Cy7 (3G8, BD-Biosciences), and anti-CD11b-APC-Cy7 (ICRF44, BD-Biosciences) to identify monocytes and separate them into activate, inflammatory, classical and non-classical subsets. Dendritic cells were identified by staining for lineage markers (CD3, CD14, CD16, CD19, PE-TXR, BD Biosciences) in a general dump gate. Lineage negative cells that were also anti-HLA DR+-APC-Cy7) (LN3, ebiosciences) (Lin-/HLA DR+) were then stained for anti-CD11c-PE-Cy7 (3.9, ebioscience), and anti-CD123-APC (7G3, BD Biosciences) to separate dendritic cells into myeloid dendritic cells (CD11c+, CD123-), and plasmacytoid cells (CD11c-, CD123+). Cells were fixed in Cytofix buffer (BD-Biosciences) and stored at -80°C in freezing medium until analyzed using flow cytometry. On the day of analysis cells were thawed, washed, and permeabilized with Cytofix/Cytoperm (BD-Biosciences) and Perm/Wash buffer (BD-Biosciences) for intracellular cytokine staining with anti-interleukin-10 (IL-10) Pacific Blue (JES3-9d7, eBioscience), anti-interleukin-12 p70 (IL-12) PE (20C2, BD - Biosciences), anti-interleukin-6 (IL-6) fluorescein isothiocyanate (MQ2-13A5, eBioscience), and anti-tumor necrosis factor-α (TNF-α) Alexafluor 700 (MAB11, BD Biosciences) for monocyte staining. Dendritic cells intracellular staining used the following antibodies: anti-IFN-α-FITC (pbl interferon source, 21112-3), anti-interleukin-12 p70 (IL-12) PE (20C2, BD -Biosciences), anti-IL-6-Alexaflour 700 (ebioscience MQ2-13A5), anti-tumor necrosis factor-α (TNF-α)-Pacific blue (ebioscience, Mab11). Samples were run using a Fortessa LSRII flow cytometer (Becton Dickinson) and analyzed using FlowJo software (FlowJo, LLC). A general gating strategy is show in Supplemental Figure 1. Cells were also analyzed for Dectin-1 surface expression (anti-Dectin-1, 15E2 Ebioscience) by flow cytometry.

### RNA seq studies/Monocyte sorting and whole glucan particle stimulation

Isolated PBMCs were washed and stained for CD14 PE-TXR and CD16 PE-Cy7 for 30 mins at 4°C in the dark. The cells were then filtered and resuspended in FACS buffer and sorted using a BD FACS Aria II. The sorted CD14+CD16+ monocytes were transferred into a round bottom 96 well plate and stimulated with whole glucan particle (10mg/ml) for 18 hours at 37°C in a 5% CO2 humidified environment. The cells were further processed for RNA isolation by using Qiagen RNeasy kit as per the manufacturer’s instructions.

### RNA-seq

The library preparation and sequencing were performed in Yale Stem Center Genomics Core Facility by Dr. Mei Zhong. The raw FASTQ files were executed for adaptor trimming and quality trimming using Cutadapt v1.18 (Martin). The quality control of the raw and trimmed FASTQ files were performed using FASTQC v0.11.7 (S., 2010). The trimmed files were aligned to human genome reference (GRCh38.primary assembly) by using STAR/ 2.7.3a-foss-2018b with parameters –runThreadN 12 --outFilterMultimapNmax 1 --outFilterMismatchNmax 999 -- outFilterMismatchNoverLmax 0.02 --alignIntronMin 20 --alignIntronMax 1000000 --alignMatesGapMax 1000000 --sjdbOverhang 100 --outSAMtype BAM SortedByCoordinate (Dobin et al., 2013). The BAM files were further sorted using SAMtools/1.9-foss-2018b and raw counts were obtained using Subread/2.0.0-GCC-7.3.0-2.30 featureCounts package and gencode v29 annotation.

### Differential gene expression

The counts from protein-coding genes, ncRNA and lincRNA was analyzed for differential gene expression in WGP Stimulated vs. Unstimulated samples using DESeq2 in R 4.0.3. Briefly, the counts were pre-filtered to remove low read (<10). For batch correction, co-variate batch was included in the design (design=∼batch+condition). The genes with a log fold change cut of 1.2 and FDR of ≤ 0.1 were considered differentially expressed genes (DEGs). The normalized counts were also obtained using DESeq2 for further analysis. The reads obtained from Y-chromosome were not included in the analysis.

### Gene set enrichment analysis (GSEA)

Gene set enrichment analysis was carried out on the normalized counts using Broad Institute GSEA v 4.1.0 (Subramanian et al., 2005). The Hallmark, C2 REACTOME, and C7 ImmuneSigDB gene sets v7.4 from Molecular signature database (MSigDB) were used for the analysis (Liberzon et al., 2011). The GSEA was run at 1000 permutation. The gene set with FDR of less than 5% was considered significant.

### Functional enrichment and Protein-Protein interaction analysis

The upregulated and downregulated DEGs were analyzed for the functional enrichment analysis by using Enrichr web server (E. Y. Chen et al., 2013; Kuleshov et al., 2016). The enrichment was performed for GO molecular function and biological processes and for Kyoto Encyclopedia of Genes and Genomes (KEGG). The significance of the enrichment was calculated using Benjamini-Hochberg FDR and terms with FDR of less than 1% were considered significant. The prediction of protein-protein interaction network was made for DEGs using searching tool for the retrieval of interacting genes (STRING) database (Szklarczyk et al., 2019). The minimum interaction score of 0.4 was used for interaction prediction. The networks of highly interacting nodes were further visualized using Cytoscape v 3.9.0 (Shannon et al., 2003). The hub genes were identified based on the degree of node using Cytohubba plugin (Chin et al., 2014).

### Upstream regulator analysis

The Qiagen Ingenuity Pathway Analysis (IPA) upstream regulator analysis (Kramer, Green, Pollard, & Tugendreich, 2014) identifies the genes that are upstream regulators and thus regulate downstream target genes. The prediction of upstream regulators was performed for the DEGs. The upstream regulators were identified based on the activation or inhibitions z-score. The z-scores with the cutoff of 2 and p-value <0.05 were considered significant.

### Statistical Analyses for Flow Cytometry studies

To assess overall cytokine expression as a function of specific participant characteristics, a series of non-parametric tests of medians were computed for each cytokine between specific subsets of the patient group (Older vs. Younger, among HIV-positive; Older vs. Younger, among HIV-negative; HIV-positive vs. HIV-negative, among Older patients; and HIV-positive vs. HIV-negative, among Younger patients). Results were FDR corrected for the set of cytokines assess within each cell type (Inflammatory monocyte, activated monocyte, classical monocyte, non-classical monocyte, and myeloid dendritic cell, plasmacytoid dendritic cells). To further explore the relationship of HIV status and age within specific cytokine expressions, parametric multivariable regression models were run for Dectin-1. First, we ran an unadjusted model that included HIV status (HIV-positive/HIV-negative), age (Older/Younger) and an HIV by age interaction. Second, we included an adjusted model that also included potentially relevant covariates. These included indicators for the use of drugs, the presence of a fungal infection, a four-level categorical variable indexing the number of comorbid conditions, and the percent of the individual’s lifespan and they have lived with HIV. Finally, we used stepwise backwards selection on the fully adjusted model to derive a parsimoniously specified model. All statistics were carried out using SAS software. Copyright, SAS Institute Inc. SAS and all other SAS Institute Inc. product or service names are registered trademarks or trademarks of SAS Institute Inc., Cary, NC, USA

### Study approval

This study was approved according to a protocol approved by the Human Research Protection Program of the Yale School of Medicine. Informed consent was obtained from all subjects prior to participation.

## Author Contributions

(Listed in order of author appearance). AK did the RNA sequencing experiments, led the analysis of the RNA-sequencing data, made all sequencing figures, and assisted in writing the manuscript. JW and HZ assisted and advised on the analysis of the RNA sequencing data. CR helped manage our database for clinical characteristics for our study participants. BVW and HA led the statistical analysis for the manuscript. ST was our data manager. LB assisted in subject recruitment efforts. SM provided vital flow cytometry training and flow cytometry analysis guidance. HZ provided expertise in bioinformatics analysis. AS provided mentorship, guidance and feedback. HJZ was the primary investigator of this study. She started the study by doing all of the flow cytometry experiments, guided the RNA seq analysis, and wrote the manuscript with AK.

## Acknowledgements

I would like to thank the subjects and staff who supported and participated in this study from the Yale Nathan Smith Clinic, Yale Health Center, and the Yale Primary care center. A special thank you goes out to the Nathan Smith clinic for their unfailing support. A special thank you to our study coordinator Allison Nelson for all her help with recruitment efforts. I would also like to thank Mei Zhong and the Yale Stm Cell Genomics Core facility for their assistance with the RNA sequencing.

