## Supplemental Files for "Dectin-1 Stimulation Promotes a Distinct Inflammatory Signature in the Setting of HIV-infection and Aging"

a.

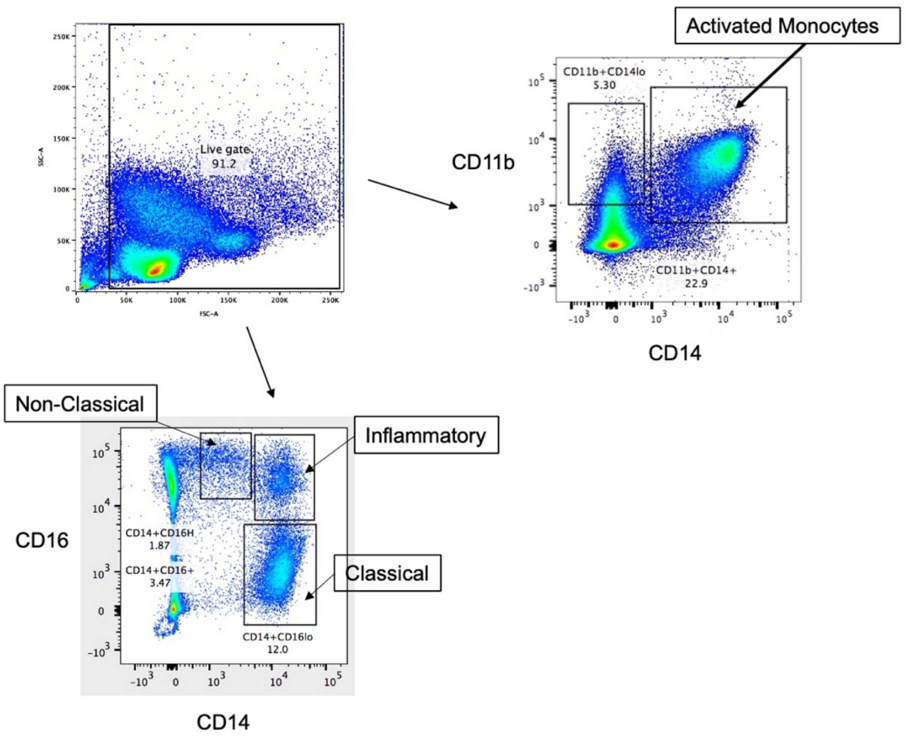

Activated Monocytes
Live gate
91.2
SSC-A
FSC-A
CD11b+CD14lo
5.30
CD11b+CD14+
22.9
CD11b
CD14
Non-Classical
Inflammatory
Classical
CD16
CD14+CD16H
1.87
CD14+CD16+
3.47
CD14+CD16lo
12.0
CD14


b.

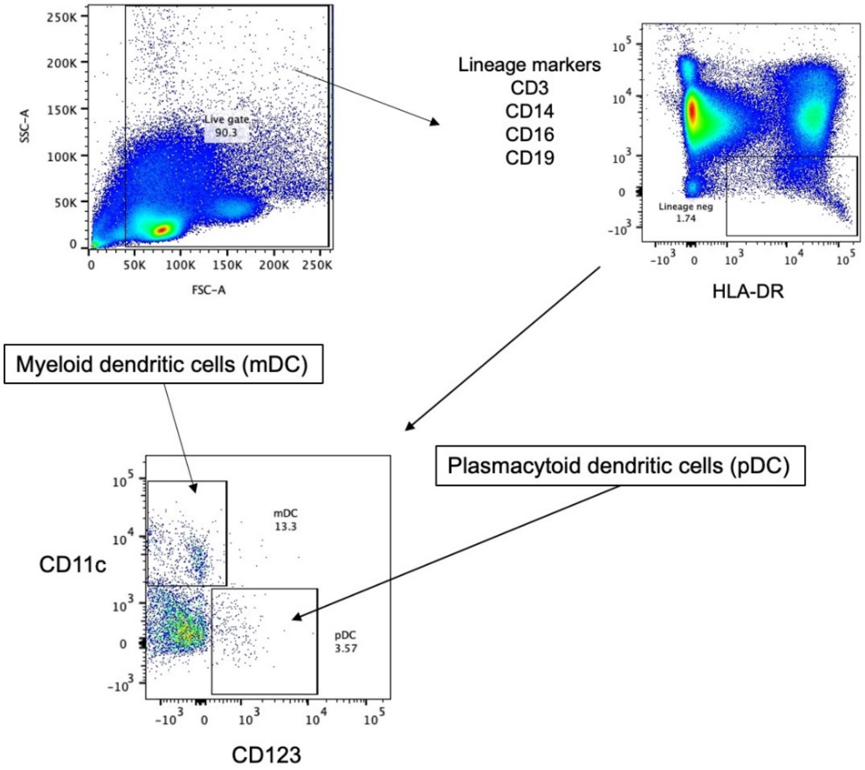

Live gate
90.3
SSC-A
FSC-A
Lineage markers
CD3
CD14
CD16
CD19
Lineage neg
1.74
HLA-DR
Myeloid dendritic cells (mDC)
Plasmacytoid dendritic cells (pDC)
mDC
13.3
pDC
3.57
CD11c
CD123

c.

| | mDC (%) | MFI IFN-α | MFI TNF-α |
|---|---|---|---|
| Baseline | 11.4 | 59.2 | -203 |
| Stimulation | 26.3 | 112 | 1828 |

| | pDC (%) | MFI IFN-α | MFI TNF-α |
|---|---|---|---|
| Baseline | 11.4 | 49.4 | -241 |
| Stimulation | 26.3 | 85.5 | 352 |

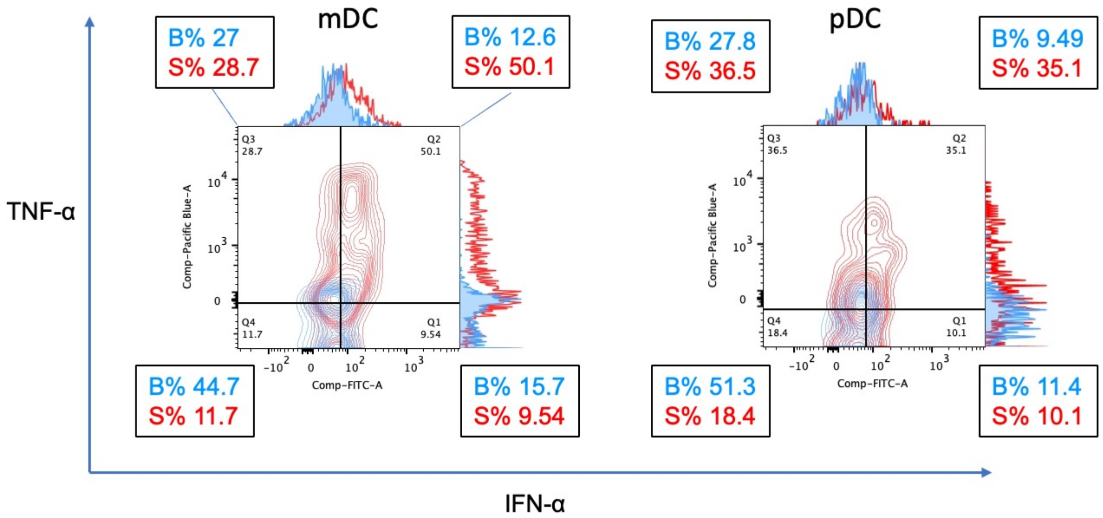


**Supplemental Figure 1, Flow Cytometry Gating Strategy and Cytokine Analysis.** a) **Monocyte Gating Strategy**. PBMCs were surface stained with anti-CD14-PE-CF594 (3G8, BD-Biosciences), anti-CD16-PE-Cy7 (3G8, BD-Biosciences), and anti-CD11b-APC-Cy7 (ICRF44, BD-Biosciences) to identify monocytes and separate them into activated (CD11b+CD14+), Inflammatory (CD14+CD16+), Classical (CD14+CD16lo), and non-classical monocytes (CD14+CD16H). Figure 1a shows the gating strategy for monocytes. All gating started with a live gate. Compensations were done for every experiment, along with isotype controls. Flow cytometry was analyzed using FloJo (LLC).

b) **Dendritic Cell Gating Strategy**. Dendritic cells were identified by staining for lineage markers (CD3, CD14, CD16, CD19, PE-TXR, BD Biosciences) in a general dump gate. Lineage negative cells that were also anti-HLA DR+-APC-Cy7) (LN3, ebiosciences) (Lin-/HLA DR+) were then stained for anti-CD11c-PE-Cy7 (3.9, ebioscience), and anti-CD123-APC (7G3, BD Biosciences) to separate dendritic cells into myeloid dendritic cells (CD11c+, CD123-), and plasmacytoid cells (CD11c-, CD123+). All gating started with a live gate. Compensations were done for every experiment, along with isotype controls. Flow cytometry was analyzed using FloJo (LLC).

c) **Cytokine Gating Strategy**. Specific cell populations (shown here are mDC and pDC populations) were then gated on to evaluate cytokines using quadrant analysis as shown in Fig 1c. Percent (%) change represents the percentage of cytokine (TNF-a, IFN-a) positive dendritic cells ( shown here are the gating for both mDCs and pDCs) that is positive for a cytokine when the baseline is subtracted from the stimulation. Quadrants were added up to get the overall % change. For example for the mDC population TNF-$\alpha$ we did the following calculation: Quadrant 3 + Quadrant 2 = % positive for TNF-$\alpha$ (27% + 12.6% = 39.6 % positive for TNF-$\alpha$ at Baseline, 28.7 + 50.1% = 78.8 % positive for TNF-$\alpha$ with Stimulation). Percent change (%) of TNF-$\alpha$ with Stimulation would be 39.2% (78.8% - 39.6%). Mean Fluorescence Intensity was also reviewed for all cytokines to make sure it correlated with dot plots and shown here with histograms and table. B% (Baseline percent positive), S% (Stimulation percent positive). Corresponding MFI values are shown in table, alongside % of mDCs and pDCs (overall of parent).

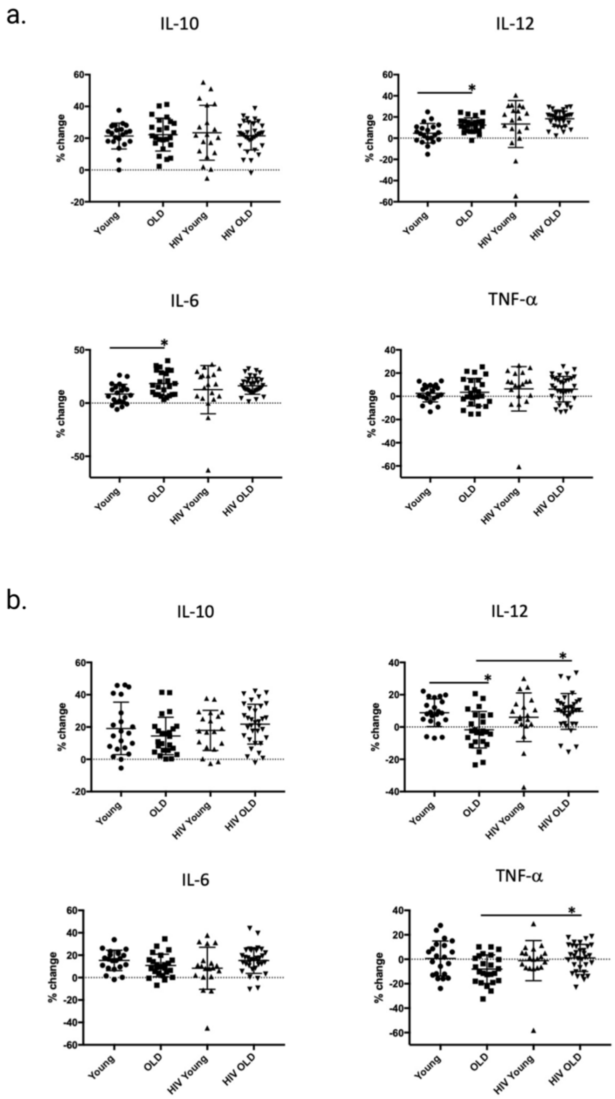


**Supplemental Figure 2**. **Effects of age and HIV-infection on Dectin-1 induced cytokine production in Classical and Non-Classical Monocytes** a) **Total cytokine production in classical monocytes (CD14+CD16lo)**. HIV-negative older adults versus young adults IL-12 (p= 0.0281), IL-6 (p= 0.0281). All other comparisons were not significant. b) **Total cytokine production in nonclassical monocytes (CD14lo, CD16H).** HIV-negative older adults versus young adults IL-12 (p= 0.044), HIV-positive older adults versus HIV-negative older adults, IL-12 (p=0.0488), TNF-α (p= 0.0488). All other comparisons were not significant.

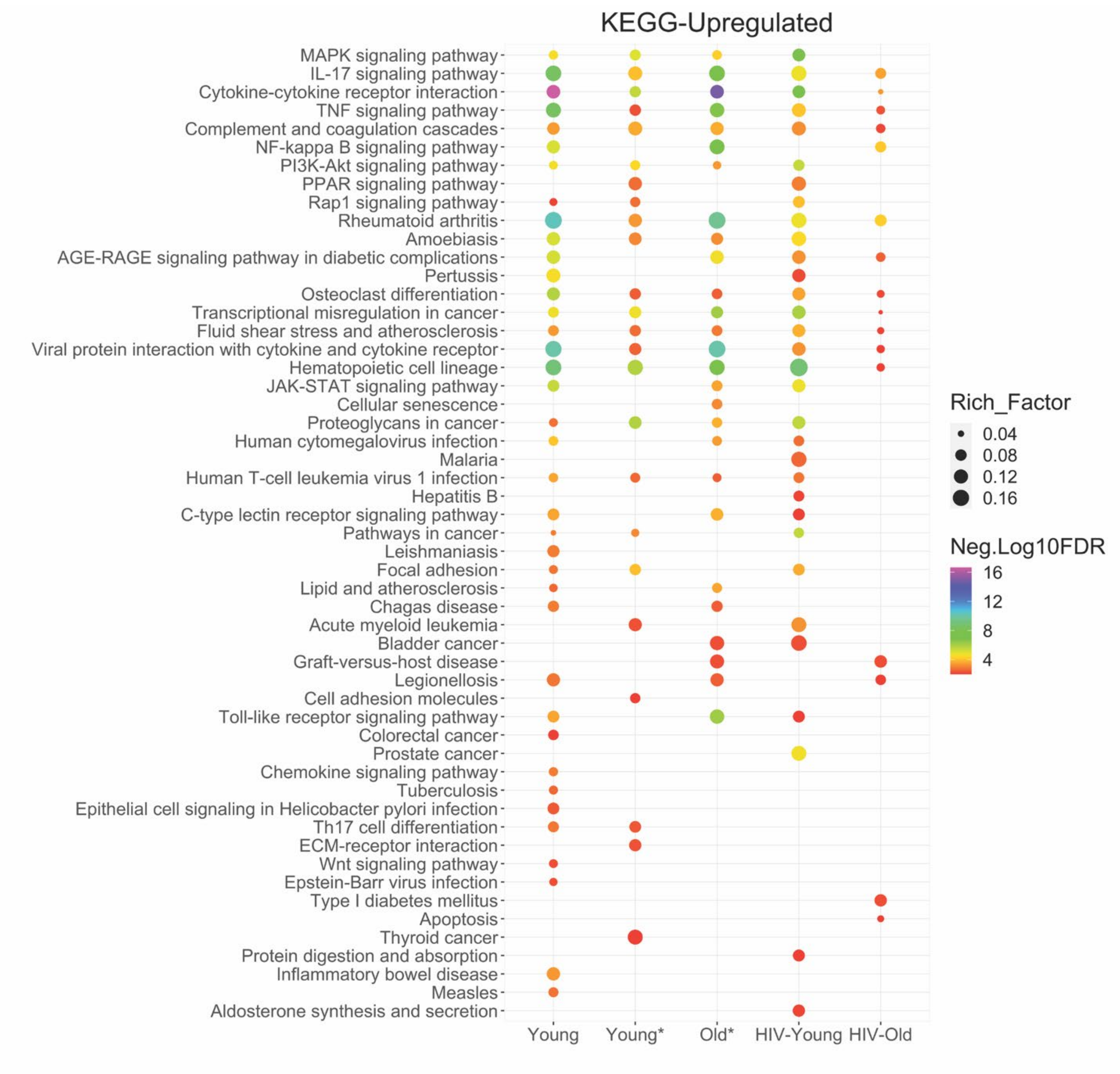


**Supplemental Figure 3**. **KEGG Pathway Analysis of Dectin-1 stimulated Monocytes**. KEGG analysis was performed for Dectin-1 stimulated Inflammatory monocytes of all cohorts using Enrichr. Dot plots represents significant KEGG pathways with FDR of <u><</u> 1%. The size of the node represents enrichment factor which is defined by overlap of the input to the gene set, color of node represents -Log10(FDR).

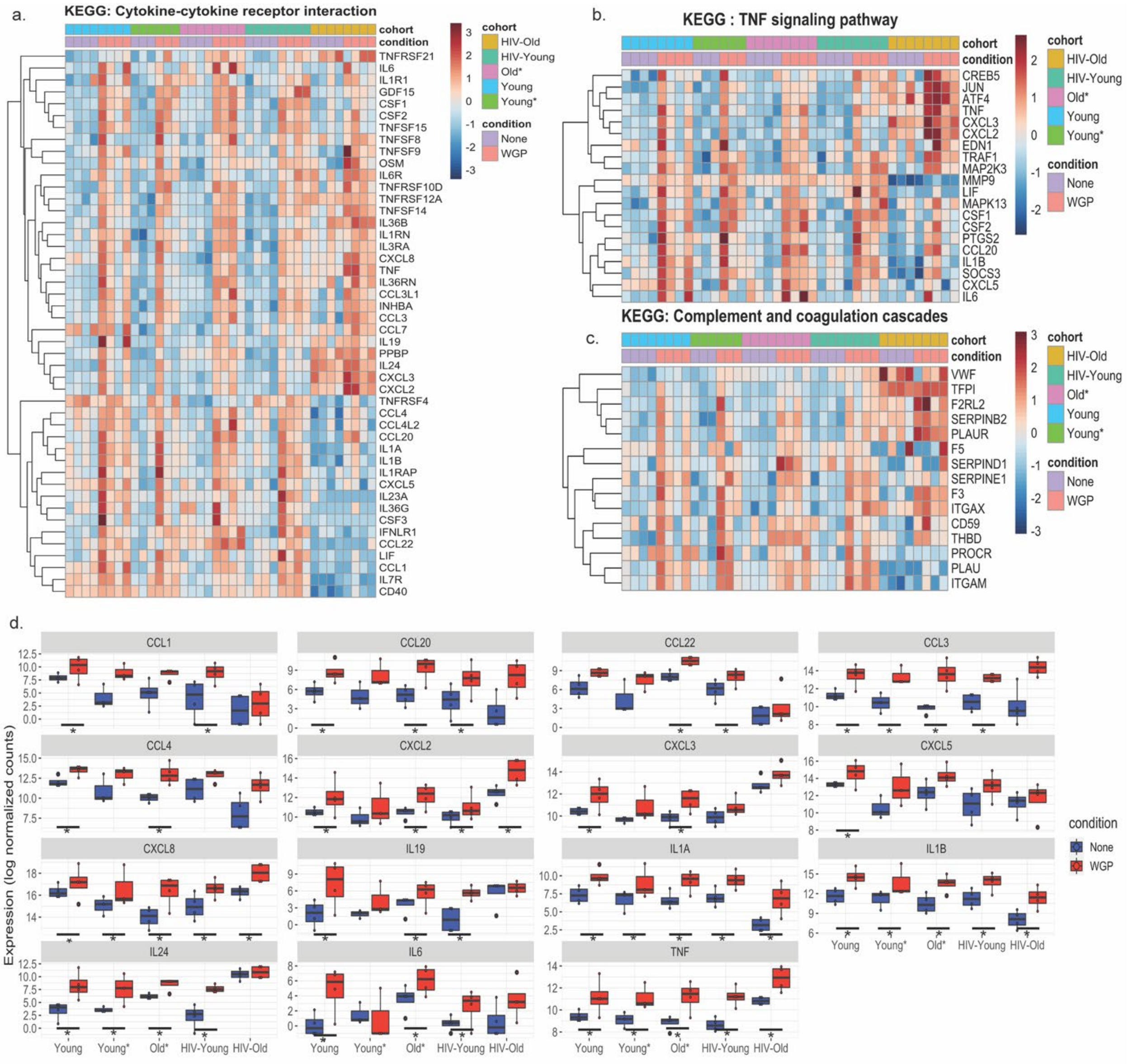


**Supplemental Figure 4**. **Dectin-1 stimulation promotes TNF-$\alpha$ signaling, and induction of the coagulation cascade in the HIV-Older cohort**. Heatmaps showing KEGG gene sets of Inflammatory Response, TNF-$\alpha$ and coagulation signaling. a) cytokine-cytokine receptor interaction and b) TNF signaling pathways and c) Complement and coagulation cascades. d) Boxplots represents log normalized counts of cytokines and chemokines in unstimulated and WGP stimulated Inflammatory monocytes. The symbol * represents significant differentially expressed genes with fold cutoff of 1.2, and q value < 0.1.

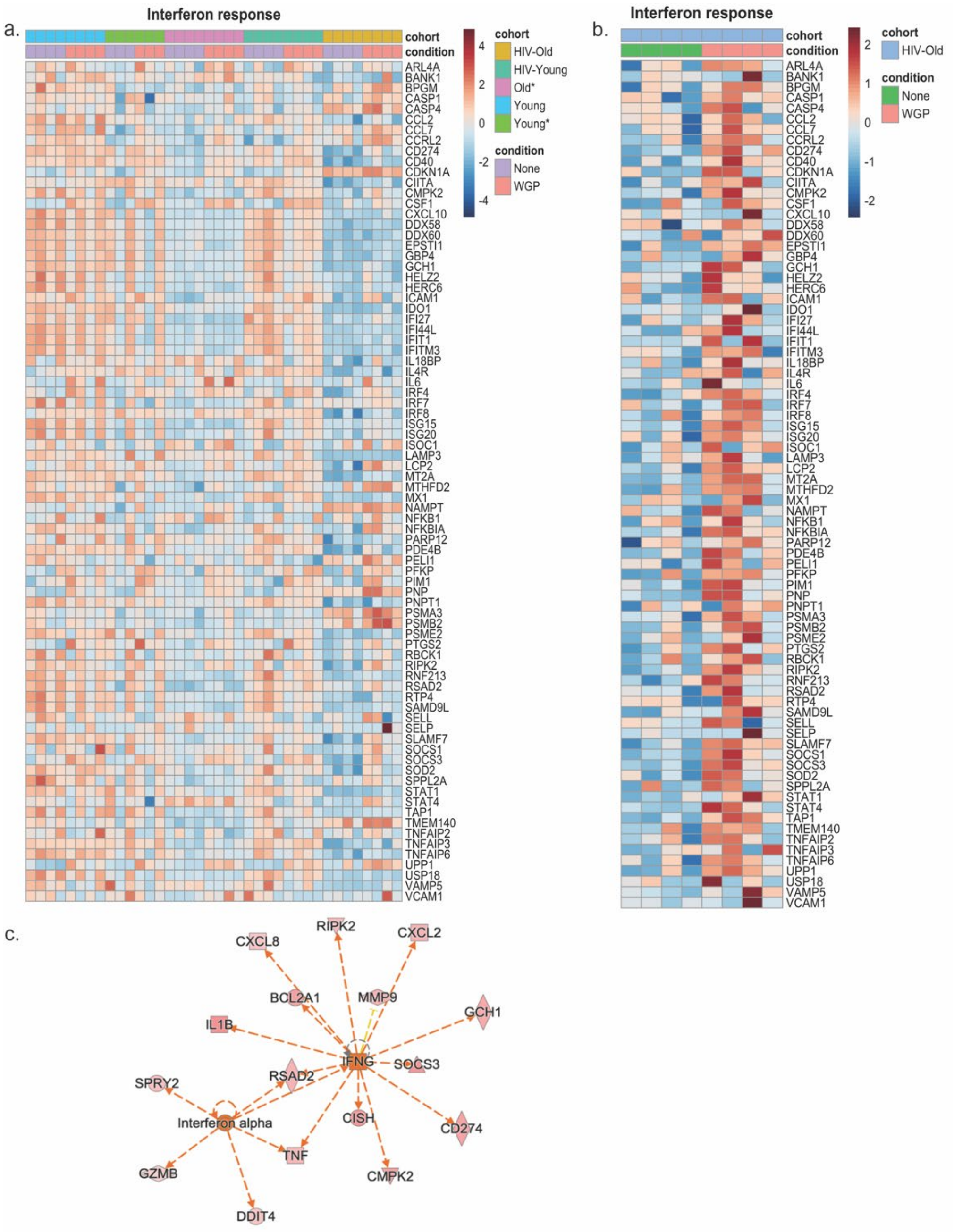


**Supplemental Figure 5. IFN-α/γ response is enriched in HIV-infected Older Adults with Dectin-1 Stimulation.** a) Heat map showing the expression profile of the gene sets that showed enrichment for Hallmark Interferon alpha and gamma response in Dectin-1 stimulated monocytes of HIV-older adults compared to other groups. b) Heatmap focusing on the response in HIV older adults, showing expression profile of the gene sets that showed enrichment for

Hallmark Interferon alpha and gamma response. c) IPA upstream analysis shows IFN-γ and IFN-α as an upstream regulators and their target genes in Dectin-1 stimulated monocytes of HIV-older adults. Volcano plot showing specific upregulated genes among HIV-Older adults in response to Dectin-1 stimulation.

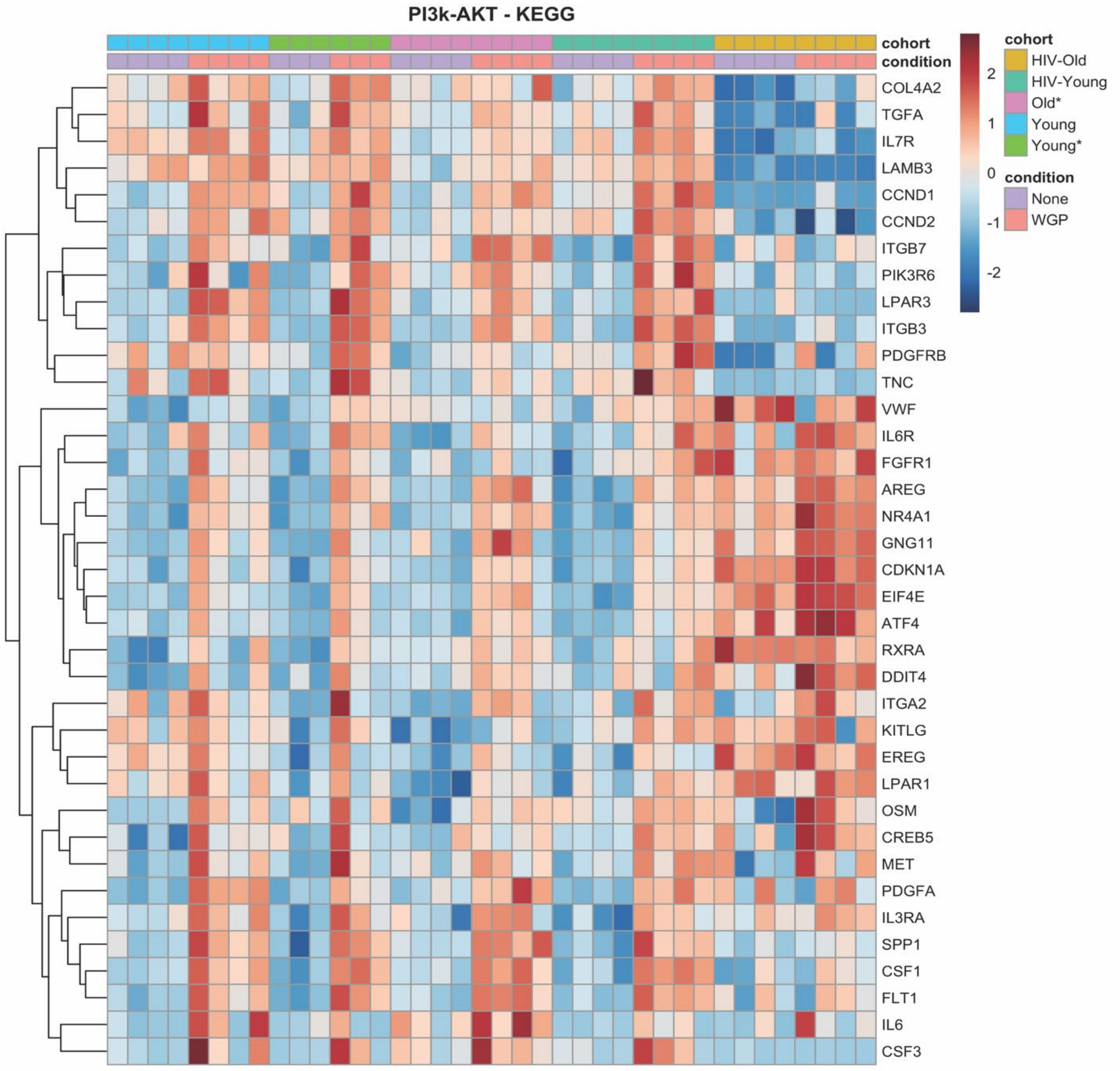


**Supplemental Figure 6. PI3K-Akt pathway is upregulated at Baseline in the HIV-Older Cohort.** Heatmap showing expression profile of genes significantly enriched for PI3k-Akt pathway during KEGG analysis. The heatmap was constructed using Pheatmap. The transcripts were normalized using variance stabilizing transformation function. The color represents relative expression of transcripts that covary across cohorts and condition.

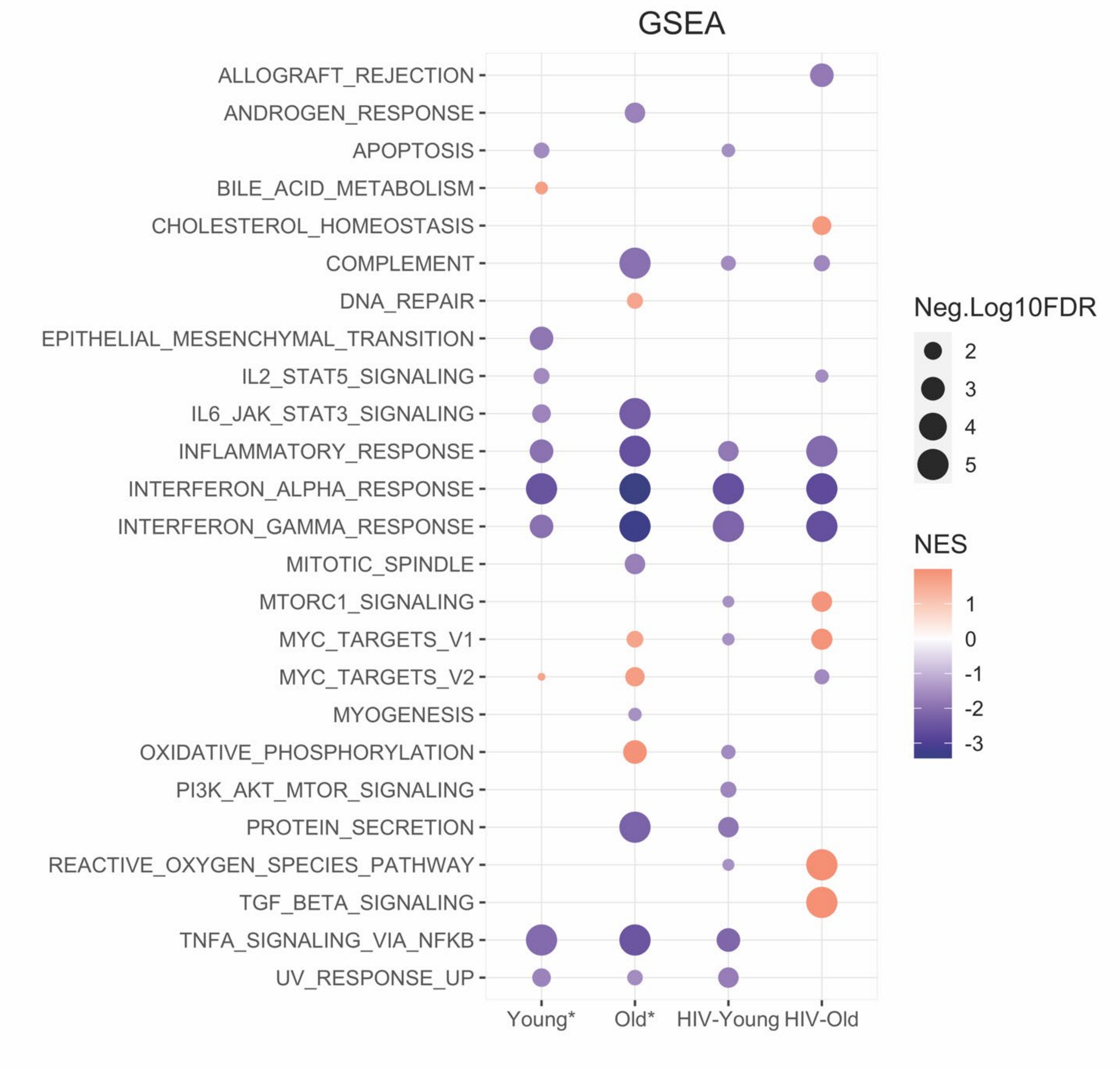


**Supplemental Figure 7. GSEA/Hallmark Baseline Analysis**. Baseline Gene Set Enrichment Analysis was performed using the Hallmark gene sets for each cohort. The dot graph represents the significant Hallmark pathways identified in the isolated monocytes of a respective cohort when compared to monocytes of healthy young individuals. The pathways with FDR of <u><5</u>% were considered significant. For graphical representation, FDR values with 0 were adjusted to 0.00001. The size of the node represents -Log10(FDR) and color of node represent normalized enrichment score. NES = Normalized enrichment scores.

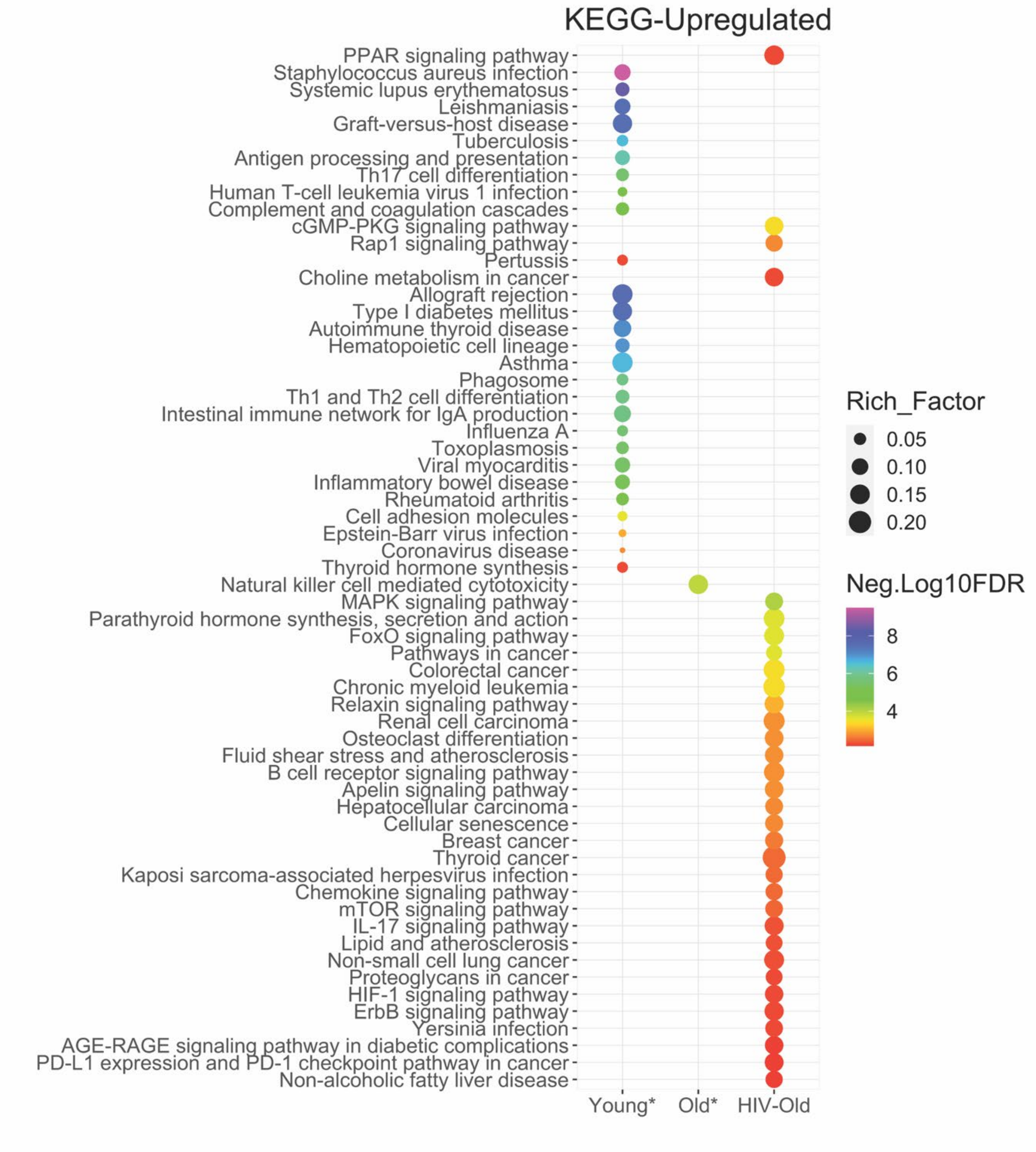


**Supplemental Figure 8. KEGG Baseline Pathway Analysis.** KEGG analysis was performed using upregulated DEGs from baseline Inflammatory monocytes of all cohorts when compared to monocytes isolated from young healthy individuals. Dot plots represents significantly upregulated KEGG pathways with FDR of $\leq 1$% compared to young healthy individuals. The size of the node represents the enrichment factor defined by overlap of the input to the gene set; color of node represents -Log10(FDR).

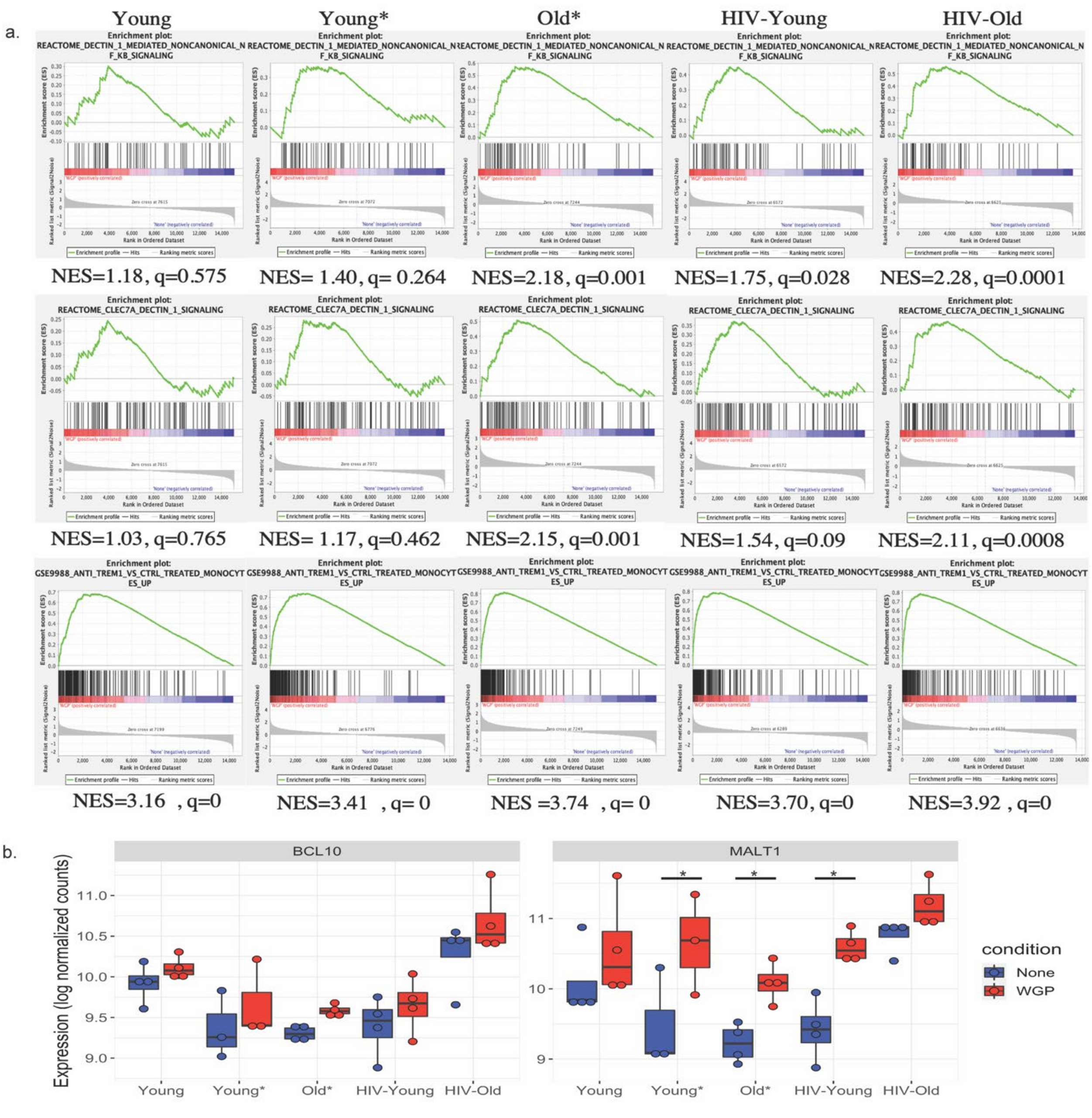


**Supplemental Figure 9. Activation of DECTIN-1/CLEC7A and TREM1 signaling in Inflammatory monocytes**. Gene set enrichment analysis (GSEA) was performed on normalized counts from all cohorts independently. Enrichment plots are shown. a) Reactome Dectin-1 mediated noncanonical NF-kB signaling, Reactome CLEC7A Dectin-1 signaling andImmuneSigDB TREM-1 signaling in monocytes. The respective normalized enrichment scores (NES) and q-values (FDR) are mentioned for each group. The scores towards the left side (red) represents the gene sets positively correlated with WGP, while scores towards the right side (blue) represents the gene sets negatively correlated with WGP. b) **Bcl10 and MALT1 are upregulated in all cohorts**. Boxplots showing log normalized counts of BCL10 and MALT1 in the unstimulated and WGP stimulated CD14+CD16+ monocytes. The symbol * represents significant differentially expressed genes with fold cutoff of 1.2, and q value < 0.1.

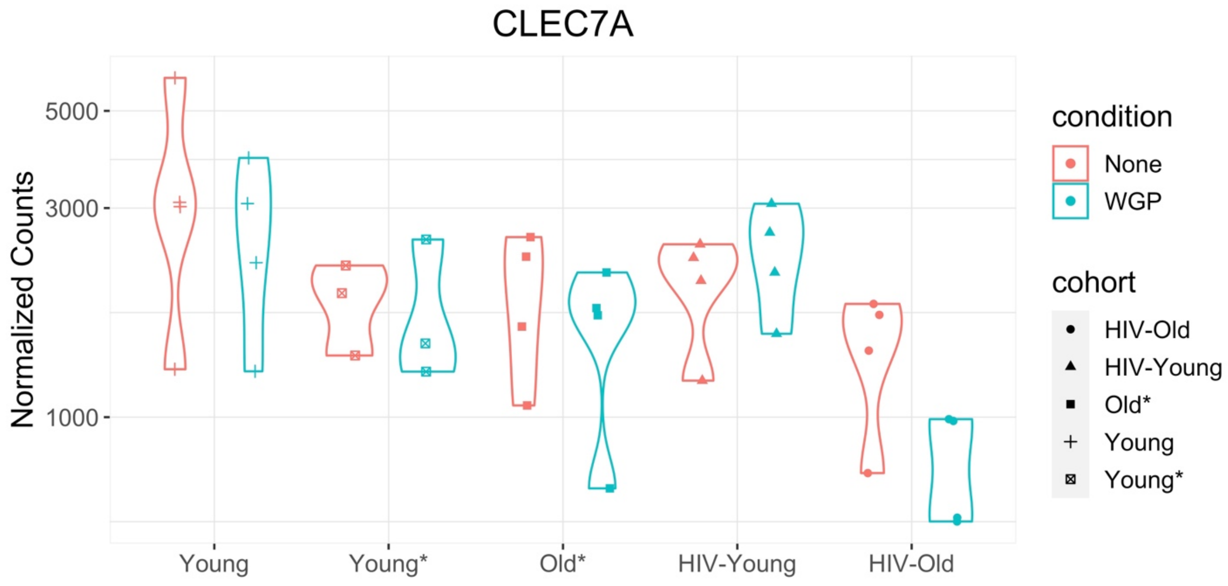


**Supplemental Figure 10. CLEC7A (Dectin-1) Expression.** Violin plot is shown representing normalized counts of CLEC7A gene expression in unstimulated and WGP stimulated CD14+CD16+ monocytes among all cohorts. The plotCounts function of DEseq2 was used to plot counts. Each point represents an individual.

**Supplemental Table 1. Complete List of Significantly Predicted Upstream Regulators.** Upstream regulator prediction was made using QIAGEN Ingenuity Pathway Analysis (IPA). DEGs were analyzed for upstream regulators using IPA. The upstream regulators with Z-score of +2 are considered as significant. The upstream regulator and their Z-scores are listed for each cohorts.

| Upstream Regulators | Young | PCC-Young | PCC-Old | HIV-Young | HIV-Old |
|---|---|---|---|---|---|
| BCR (complex) | 3.1 | 2.219 | 3.257 | 2.952 | N/A |
| MAPK1 | 1.757 | 2.213 | 1.876 | 5.383 | N/A |
| IL15 | 2.442 | 2.778 | 2.414 | 3.256 | N/A |
| ASPSCR1-TFE3 | 2.236 | 3 | 2.449 | 3 | N/A |
| MMP1 | 2.4 | 2.598 | 2.598 | 2.954 | N/A |
| ADORA3 | 2.646 | 2.449 | 3 | 1.89 | N/A |
| KRAS | 2.449 | 2.449 | 2.236 | 2.772 | N/A |
| EZH2 | 3.293 | 1.34 | 2.834 | 2.424 | N/A |
| IL5 | 2.219 | 2.433 | 2.433 | 2.621 | N/A |
| let-7 | -2.066 | -2.643 | -2.25 | -2.59 | N/A |
| COL18A1 | -2.897 | -2.354 | -2.36 | -1.884 | N/A |
| PDCD4 | -2.621 | -1.982 | -2.425 | -2.425 | N/A |
| mir-8 | -2.236 | -1.987 | -2.438 | -2.407 | N/A |
| PAF1 | 2.433 | 2.236 | 2.63 | -1.698 | N/A |
| STAT5a/b | 2.219 | 2.425 | 2.213 | 1.937 | N/A |
| CRP | 2.191 | 1.995 | 2.191 | 2.412 | N/A |
| miR-146a-5p (and other miRNAs w/seed GAGAACU) | -1.664 | -2.433 | -2.219 | -2.433 | N/A |
| SP1 | 2.157 | 1.976 | 2.767 | 1.747 | N/A |
| PI3K (complex) | 2.371 | 1.746 | 2.573 | 1.926 | N/A |
| ERG | 1.914 | 2.339 | 2.186 | 2.037 | N/A |
| BRD4 | 2.345 | 2.561 | 2.557 | 0.98 | N/A |
| CBX5 | -2.121 | -2.646 | -2.236 | -1.387 | N/A |
| ATF4 | 1.98 | 1.478 | 2.391 | 2.432 | N/A |
| RAC1 | 2.236 | 1.89 | 1.89 | 2.121 | N/A |
| ETV1 | 2 | 2 | 2 | 2 | N/A |
| RNASE1 | 2.63 | 2.236 | 2.813 | -0.128 | N/A |
| RNASE2 | 2.449 | 2.204 | 2.779 | 0.286 | N/A |
| VEGFA | 2.383 | 1.679 | 1.764 | 1.871 | N/A |
| FOXM1 | 2.726 | 1.033 | 1.933 | 1.994 | N/A |

| | | | | | |
|---|---|---|---|---|---|
| ATM | 1.284 | 1.657 | 2.568 | 1.865 | N/A |
| SAA | 2.391 | 2.391 | N/A | 2.391 | N/A |
| Lymphotoxin | 2.449 | N/A | 2.449 | 2.236 | N/A |
| CEBPB | 2.165 | 1.183 | 2.619 | 1.065 | N/A |
| NR5A2 | 2.234 | 1.506 | 1.506 | 1.765 | N/A |
| PTAFR | 1.997 | 0.905 | 2.218 | 1.831 | N/A |
| TLR9 | 1.564 | 0.806 | 3.065 | -1.487 | N/A |
| CIC | N/A | -2.449 | -2.236 | -2.236 | N/A |
| SREBF1 | 1.934 | 1.346 | 2.166 | 1.46 | N/A |
| NEDD9 | 2 | 1.195 | 2.236 | 1.463 | N/A |
| FCER1G | 2.219 | N/A | 2.433 | 2.236 | N/A |
| PDPK1 | -2.198 | N/A | -2.408 | -2.225 | N/A |
| FCGR2A | 2.225 | N/A | 2.411 | 2.195 | N/A |
| ETS1 | 2.22 | 2.63 | N/A | 1.968 | N/A |
| NORAD | N/A | 2.236 | 2.219 | 2.345 | N/A |
| PIN1 | 2.191 | N/A | 2.201 | 2.408 | N/A |
| SIRT6 | 2.19 | N/A | 2.395 | 2.19 | N/A |
| IFNL1 | N/A | N/A | 1.94 | -4.821 | N/A |
| CREB1 | 2.42 | N/A | 2.041 | 2.224 | N/A |
| CD14 | 2.215 | N/A | 2.431 | 1.981 | N/A |
| TLR6 | 1.982 | N/A | 2.425 | 2.213 | N/A |
| Lh | 2.195 | 0.878 | 2.606 | 0.94 | N/A |
| TERT | N/A | 1.98 | 2.425 | 2.213 | N/A |
| Nr1h | -2.044 | N/A | -2.276 | -2.236 | N/A |
| IKBKB | 1.963 | N/A | 2.6 | 1.981 | N/A |
| IL1 | 1.982 | N/A | 2.219 | 2.219 | N/A |
| TRAF6 | 1.982 | N/A | 2.433 | 2 | N/A |
| PLAUR | 1.981 | 2.207 | N/A | 2.207 | N/A |
| MAPK8 | 2.2 | 1.423 | 1.095 | 1.664 | N/A |
| PDLIM2 | 2.121 | 1.134 | 1.897 | 1.213 | N/A |
| MAPK14 | 2.2 | N/A | 2.2 | 1.964 | N/A |
| CHUK | 2.213 | N/A | 2.4 | 1.746 | N/A |
| TLR5 | 1.966 | N/A | 2.407 | 1.98 | N/A |
| TLR1 | 1.969 | N/A | 2.401 | 1.972 | N/A |
| EHF | 2.449 | 1.265 | 1.89 | 0.707 | N/A |
| FIRRE | 2 | N/A | 2 | 2.236 | N/A |
| FASN | -1.964 | N/A | -2.236 | -2 | N/A |
| PCDH11Y | 1.982 | 1.982 | N/A | 2.213 | N/A |
| FAT1 | 2.213 | N/A | 1.974 | 1.974 | N/A |
| THRB | -1.455 | -1.501 | -2 | -1.2 | N/A |
| KDM3A | N/A | 2.19 | 1.96 | 1.969 | N/A |
| APP | 2.4 | N/A | 2.4 | 1.303 | N/A |
| EDN1 | 1.954 | N/A | 2.183 | 1.954 | N/A |
| MACROH2A1 | -2.121 | -0.816 | -2.333 | -0.707 | N/A |
| CRNDE | 2.236 | 1.969 | 0.509 | 1.242 | N/A |

| | | | | | |
|---|---|---|---|---|---|
| EHMT1 | 2.236 | 2.111 | 0.447 | 1.155 | N/A |
| mir-146 | -1.96 | N/A | -2.407 | -1.452 | N/A |
| MUC1 | 1.785 | 2.176 | 0.584 | 1.22 | N/A |
| FSH | 1.989 | 0.626 | 2.434 | 0.698 | N/A |
| IL33 | 2.213 | 0.689 | 1.392 | 1.443 | N/A |
| ABL1 | N/A | 1.951 | 2.186 | 1.406 | N/A |
| TFRC | -2.4 | -1.23 | N/A | -1.723 | N/A |
| P2RX7 | 2.2 | N/A | 1.96 | 1.131 | N/A |
| EFNA1 | -1 | -1.89 | N/A | -2.333 | N/A |
| RBPJ | N/A | -2.646 | N/A | -2.449 | N/A |
| EFNA2 | -1 | -1.89 | N/A | -2.121 | N/A |
| MAFB | N/A | -1.89 | N/A | -3.051 | N/A |
| SRC | 1.982 | 0.13 | 2.156 | 0.656 | N/A |
| WNT5A | 1.308 | N/A | 2.211 | 1.34 | N/A |
| ESRRA | 2.421 | N/A | N/A | 2.421 | N/A |
| E2F1 | N/A | N/A | 2.415 | 2.415 | N/A |
| IRF4 | -1.492 | -2 | -0.213 | 0.784 | N/A |
| PRKAA1 | 1.067 | N/A | 1.387 | 2 | N/A |
| PRKCE | 2.164 | N/A | 0.964 | 1.304 | N/A |
| JAK1 | 2 | N/A | 2 | 0.378 | N/A |
| IRGM | N/A | N/A | N/A | 4.345 | N/A |
| SOX11 | N/A | 1.912 | N/A | 2.331 | N/A |
| PGF | N/A | N/A | 2.229 | 1.994 | N/A |
| IFIH1 | 1.987 | N/A | 2.216 | N/A | N/A |
| NR1H4 | -1.964 | N/A | -2.2 | N/A | N/A |
| ELAVL1 | N/A | N/A | 2.401 | 1.747 | N/A |
| BCL2L1 | -2.219 | -1 | N/A | -0.896 | N/A |
| SCD | N/A | N/A | -2.157 | -1.934 | N/A |
| RNY3 | N/A | N/A | N/A | -4.025 | N/A |
| HDAC1 | N/A | -2 | N/A | -2 | N/A |
| miR-182-5p (and other miRNAs w/seed UUGGCAA) | N/A | N/A | N/A | 3.86 | N/A |
| KMT2D | N/A | 2.224 | N/A | 1.59 | N/A |
| CBFA2T3 | 0.659 | 0.659 | N/A | 2.484 | N/A |
| miR-450a-5p (and other miRNAs w/seed UUUGCGA) | N/A | -2.236 | N/A | -1.414 | N/A |
| EZR | N/A | N/A | -2 | -1.633 | N/A |
| TLR8 | N/A | N/A | 2.422 | 1.177 | N/A |
| IL1RAP | N/A | N/A | 1.342 | 2 | N/A |
| RC3H1 | N/A | N/A | N/A | 3.207 | N/A |
| IFNB1 | N/A | N/A | N/A | -3.144 | N/A |
| CGAS | N/A | N/A | 2.213 | 0.796 | N/A |
| IFNL4 | N/A | N/A | N/A | -2.975 | N/A |
| IFNAR2 | N/A | N/A | N/A | -2.646 | N/A |
| JAK | N/A | N/A | N/A | -2.646 | N/A |

| | | | | | |
|---|---|---|---|---|---|
| SOCS1 | 0.192 | N/A | 0.192 | 2.177 | N/A |
| miR-122-5p (miRNAs w/seed GGAGUGU) | N/A | -2.449 | N/A | N/A | N/A |
| TSLP | N/A | N/A | 2.425 | N/A | N/A |
| IRF1 | N/A | N/A | N/A | -2.39 | N/A |
| miR-511-5p (miRNAs w/seed UGUCUUU) | N/A | N/A | -2.236 | N/A | N/A |
| ANLN | N/A | N/A | N/A | -2.236 | N/A |
| AURK | N/A | N/A | N/A | -2.219 | N/A |
| HDAC4 | N/A | N/A | N/A | -2.219 | N/A |
| miR-296-5p (miRNAs w/seed GGGCCCC) | N/A | -2.219 | N/A | N/A | N/A |
| GATA6 | N/A | N/A | N/A | 2.213 | N/A |
| IRF7 | N/A | N/A | N/A | -2.202 | N/A |
| ISG15 | N/A | N/A | N/A | 2.195 | N/A |
| IKZF3 | N/A | N/A | N/A | 2.137 | N/A |
| TAP1 | N/A | N/A | -2 | 0.112 | N/A |
| HMGB1 | N/A | N/A | 2.085 | N/A | N/A |
| PAEP | N/A | N/A | -2 | N/A | N/A |
| RUNX1 | 2 | N/A | N/A | N/A | N/A |
| WNT7A | N/A | N/A | N/A | 2 | N/A |
| NFIX | N/A | N/A | N/A | -2 | N/A |
| FCER2 | N/A | N/A | 2 | N/A | N/A |
| FENDRR | 2 | N/A | N/A | N/A | N/A |
| CLEC4E | N/A | N/A | 2 | N/A | N/A |
| LGALS8 | N/A | N/A | 2 | N/A | N/A |
| TPR | N/A | N/A | 2 | N/A | N/A |
| MAP2K3 | N/A | N/A | N/A | -2 | N/A |
| TREM1 | 7.147 | 6.929 | 7.484 | 7.617 | 5.785 |
| TNF | 5.777 | 4.211 | 6.326 | 3.016 | 4.193 |
| SMARCA4 | 5.043 | 4.483 | 4.745 | 3.872 | 4.123 |
| IL1B | 4.699 | 3.941 | 5.19 | 3.52 | 3.639 |
| RELA | 3.73 | 1.992 | 4.507 | 1.252 | 3.582 |
| FOXO1 | 2.926 | 2.596 | 2.486 | 1.86 | 3.257 |
| IL1A | 4.227 | 2.725 | 4.423 | 3.152 | 3.2 |
| IL17A | 3.89 | 2.395 | 3.872 | 2.847 | 3.197 |
| NFkB (complex) | 4.457 | 2.855 | 5.05 | 2.086 | 3.196 |
| PDGF BB | 4.783 | 4.125 | 4.911 | 4.794 | 3.162 |
| F7 | 3.13 | 2.449 | 3.108 | 2.931 | 2.945 |
| PRKCD | 3.199 | 2.944 | 3.483 | 3.281 | 2.92 |
| IFNG | 2.189 | 0.321 | 3.235 | -3.362 | 2.905 |
| JUN | 3.53 | 3.09 | 4.175 | 4.027 | 2.786 |
| OSCAR | 3.742 | 3.317 | 3.742 | 3.606 | 2.646 |
| CCL5 | 2.828 | 3.317 | 3.606 | 2.84 | 2.646 |
| SYVN1 | 3 | 2.53 | 3.317 | 2 | 2.646 |
| CHD1 | 3.464 | N/A | 3.317 | 2.646 | 2.646 |

| | | | | | |
|---|---|---|---|---|---|
| TGFB1 | 3.653 | 3.519 | 3.652 | 3.084 | 2.618 |
| CD28 | 2.965 | 1.916 | 2.589 | 2.402 | 2.594 |
| P38 MAPK | 3.178 | 2.576 | 4.007 | 1.276 | 2.586 |
| TLR7 | 2.15 | 0.265 | 3.775 | -1.094 | 2.57 |
| EGF | 2.492 | 2.819 | 3.522 | 3.303 | 2.564 |
| FOXO3 | 1.377 | 0.911 | 2.169 | 0.161 | 2.538 |
| GPER1 | 3.286 | 3.13 | 3.973 | 3.576 | 2.449 |
| Collagen type II | 3.606 | 3.162 | 3.317 | 3.317 | 2.449 |
| RNF138 | 2.646 | N/A | 2.828 | 2 | 2.449 |
| CSF2 | 4.262 | 3.885 | 4.157 | 4.123 | 2.433 |
| FOXL2 | 3.293 | 2.635 | 2.885 | 3.145 | 2.429 |
| CD36 | 3.845 | 3.162 | 3.576 | 3.01 | 2.414 |
| Immunoglobulin | 2.534 | 1.778 | 3.434 | 0.671 | 2.414 |
| FOS | 1.718 | 1.414 | 2.714 | 2.496 | 2.414 |
| CD40LG | 2.61 | 1.007 | 3.794 | 2.007 | 2.406 |
| S100A8 | 2.572 | N/A | 2.921 | 1.985 | 2.388 |
| ERK1/2 | 3.198 | 0.996 | 3.507 | 2.197 | 2.387 |
| MAP2K1/2 | 2.739 | 2.92 | 3.36 | 3.361 | 2.38 |
| HIF1A | 2.538 | 0.331 | 3.451 | 1.215 | 2.356 |
| TGM2 | 3.576 | 2.323 | 2.891 | -0.561 | 2.345 |
| TLR4 | 2.869 | 2.547 | 3.166 | 1.975 | 2.339 |
| NUPR1 | 3.273 | 3.656 | 4.271 | 4.061 | 2.333 |
| STAT3 | 3.02 | 1.265 | 2.784 | 2.023 | 2.314 |
| TLR7/8 | 2.646 | 2.449 | 3.317 | 1.897 | 2.236 |
| MAP2K1 | 2.159 | 2.139 | 2.271 | 2.35 | 2.236 |
| SELPLG | 2.449 | 1.342 | 3 | 1.414 | 2.236 |
| MTOR | 1.913 | N/A | 1.238 | 0.798 | 2.236 |
| CD3 | 2.676 | 1.706 | 2.751 | 0.444 | 2.224 |
| SMAD4 | 1.551 | 1.303 | 1.57 | 1.805 | 2.224 |
| Interferon alpha | 1.712 | 1.731 | 0.518 | -3.088 | 2.219 |
| CAMP | 3.554 | 3.252 | 3.157 | 3.667 | 2.21 |
| PPRC1 | 3.111 | 2 | 2.771 | 3.266 | 2.197 |
| NCR2 | 2.779 | 2.186 | 2.95 | 2.4 | 2.19 |
| F2 | 2.215 | 1.706 | 2.752 | 1.522 | 2.177 |
| IL18 | 2.928 | 1.999 | 3.244 | 1.329 | 2.176 |
| EGFR | 2.986 | 3.111 | 3.005 | 3.279 | 2.17 |
| TLR2 | 3.212 | 3.061 | 3.338 | 3.091 | 2.156 |
| NFKB1 | 1.982 | 1.342 | 2.887 | 0.842 | 2.129 |
| CG | 1.978 | 1.348 | 3.104 | 0.612 | 2.023 |
| JUNB | 2.121 | 2.121 | 3 | 2.714 | 2 |
| TNFSF14 | 2.643 | 2.234 | 2.612 | 1.349 | 1.999 |
| PTGS2 | 2.229 | 1.778 | 2.42 | 1.603 | 1.992 |
| PLG | 2.423 | N/A | 2.614 | 2.207 | 1.991 |
| C5 | 2.109 | 1.857 | 2.438 | 2.332 | 1.987 |
| Cdk | 2.224 | N/A | 1.854 | 2 | 1.987 |

| | | | | | |
|---|---|---|---|---|---|
| NAMPT | 1.991 | 1.274 | 2.071 | 1.612 | 1.986 |
| CD3 group | 1.953 | 2.19 | 2.2 | 2.2 | 1.982 |
| CTNNB1 | 0.935 | 1.133 | 0.369 | 2.416 | 1.982 |
| Ap1 | 2.573 | 2.804 | 2.755 | 3.132 | 1.98 |
| ERK | 4.308 | 3.528 | 3.544 | 3.815 | 1.979 |
| PF4 | 2.309 | 0.522 | 2.992 | 1.634 | 1.963 |
| S100A9 | 2.374 | 1.934 | 2.576 | 2.377 | 1.962 |
| NFAT5 | 1.961 | 1.948 | 2.395 | 1.948 | 1.961 |
| TLR3 | 2.199 | N/A | 2.406 | -0.085 | 1.955 |
| GLI1 | 2.401 | 2.719 | 1.82 | 2.37 | 1.951 |
| MYD88 | N/A | N/A | 2.189 | 1.56 | 1.951 |
| EGR1 | 2.585 | 3.082 | 2.926 | 3.641 | 1.941 |
| FGF2 | 2.449 | 1.808 | 2.449 | 1.947 | 1.941 |
| ECSIT | 2.772 | N/A | 3.264 | 2.588 | 1.941 |
| BSG | 2.412 | 1.939 | 2.607 | 1.807 | 1.934 |
| Tlr | 1.93 | N/A | 2.152 | 1.929 | 1.929 |
| Fcer1 | 2.934 | 2.401 | 3.253 | 2.568 | 1.917 |
| PRL | 1.957 | N/A | 1.489 | -4.321 | 1.914 |
| PPARD | 3.317 | 2.236 | 2.704 | 2.646 | 1.913 |
| Akt | 1.995 | 1.227 | 2.497 | 1.631 | 1.896 |
| Mek | 2.683 | 2.112 | 3.113 | 1.504 | 1.856 |
| SPI1 | 0.555 | 1.206 | 1.151 | -3.358 | 1.732 |
| Jnk | 3.051 | 1.859 | 4.16 | 2.777 | 1.706 |
| TCR | 2.187 | 2.591 | 1.038 | 2.939 | 1.698 |
| IL2 | 1.593 | 2.157 | 1.62 | 2.739 | 1.514 |
| TP63 | 2.586 | 1.423 | 2.322 | 1.475 | 1.488 |
| ERBB2 | 3.058 | 3.482 | 3.234 | 2.87 | 1.405 |
| IFNA2 | N/A | N/A | N/A | -4.266 | 1.402 |
| IGF1 | 1.006 | 1.643 | 2.553 | 1.902 | 1.308 |
| IKBKE | 2.224 | 2.229 | 1.992 | 2.49 | 1.131 |
| CD40 | 1.948 | 2.256 | 2.825 | 2.362 | 1.066 |
| DDX58 | 2.414 | N/A | 2.056 | 1.44 | 0.958 |
| BTK | 1.47 | 1.616 | 0.488 | 3.983 | 0.818 |
| HGF | 2.416 | 0.286 | 2.313 | 1.352 | 0.804 |
| PGR | 1.167 | 1.906 | 1.074 | 3.741 | 0.726 |
| STAT1 | N/A | N/A | 1.737 | -3.711 | 0.718 |
| IL27 | 1.747 | 2.136 | 1.9 | -0.421 | 0.552 |
| IL4 | 2.738 | 3.326 | 3.315 | 3.573 | 0.468 |
| estrogen receptor | -2.284 | -1.888 | -1.119 | -2.049 | 0.447 |
| ESR1 | 1.429 | 1.337 | -0.056 | 2.272 | -0.404 |
| MYC | 0.897 | -0.422 | -0.471 | 2.505 | -0.529 |
| IL13 | 1.527 | 3.353 | 1.141 | 3.299 | -0.822 |
| IL10 | -0.932 | -1.253 | -1.745 | -2.316 | -0.937 |
| NEUROG1 | -1 | -1.508 | -2.449 | -0.832 | -1 |
| GSTO1 | -1.3 | -2.086 | -0.539 | -0.184 | -1.069 |

| | | | | | |
|---|---|---|---|---|---|
| Hdac | -1.873 | -2.097 | -2.524 | -2.023 | -1.564 |
| NR3C1 | -0.75 | -1.4 | -2.597 | -1.509 | -1.782 |
| IL37 | -2.18 | N/A | -2.38 | -2.378 | -1.944 |
| HLX | -0.849 | N/A | -2.213 | -0.762 | -1.982 |
| JAG2 | -2.915 | -2.433 | -2.395 | -1.964 | -2 |
| OGA | 0 | -1.36 | -2.359 | -1.422 | -2 |
| TAB1 | -1.982 | N/A | -1.387 | 0.707 | -2 |
| WBP2 | -1 | N/A | -1.897 | -0.277 | -2 |
| COPA | N/A | N/A | -2 | -1 | -2 |
| S100A6 | N/A | N/A | -1.89 | -1 | -2 |
| PDCD1 | N/A | -0.447 | -1.342 | -0.707 | -2 |
| ARID1A | N/A | N/A | -1.166 | -0.868 | -2 |
| CIP2A | -1.387 | -0.478 | -2.433 | -1.066 | -2.219 |
| miR-155-5p (miRNAs w/seed UAAUGCU) | -3.539 | -2.789 | -3.4 | -3.095 | -2.385 |
| IL1RN | -2.177 | -0.651 | -2.371 | 2.155 | -2.449 |
| ETV6-RUNX1 | -3 | -2.692 | -3.308 | -2.031 | -2.828 |

| | HIV negative (n =11) | HIV positive (n = 8) |
|---|---|---|
| **Age Group** | | |
| 21-40 (n, %) | 7 (64%) | 4 (50%) |
| 60 or older (n, %) | 4 (36%) | 4 (50%) |
| **Female (n, %)** | 6 (55%) | 3 (38%) |
| **Race** | | |
| Black (n, %) | 5 (45%) | 4 (50%) |
| White (n, %) | 5 (45%) | 2 (25%) |
| Other (n, %) | 1 (9%) | 2 (25%) |
| **Latinx (n, %)** | 3 (27%) | 3 (38%) |
| **BMI (mean, range)** | 30.0 (22.5-41.0) | 27.9 (15.1-44.7) |
| **BMI ≥30 kg/m$^2$ (n, %)** | 4 (36%) | 3 (38%) |
| **DM2 (n, %)** | 4 (36%) | 2 (25%) |
| **Metabolic syndrome (DM2 + HTN + HLD) (n, %)** | 3 (27%) | 2 (25%) |
| **CAD and/or PVD (n, %)** | 0 | 2 (25%) |
| **Asthma and/or COPD (n, %)** | 4 (36%) | 4 (50%) |
| **Autoimmune and/or autoinflammatory disease (n, %)** | 1 (9%) | 0 |
| **Prior history of smoking (n, %)** | 2 (18%) | 2 (25%) |
| **Current smoker (n, %)** | 4 (36%) | 2 (25%) |
| **Recreational drug use in past year (n, %)** | 3 (27%) | 2 (25%) |
| **History of positive PPD or QuantiFERON (n, %)** | 0 | 2 (25%) |
| **History of fungal disease (n, %)** | 1 (9%) | 3 (38%) |
| **CD4 >200 (n, %)** | NA | 8 (100%) |
| **VL >100 (n, %)** | NA | 1 (13%) |
| **On HAART (n, %)** | NA | 8 (100%) |
| **% life span with HIV (mean, range)** | NA | 54% (14-100%) |
| **Congenital HIV (n, %)** | NA | 2 (25%) |

**Supplemental Table 2. Clinical characteristics of RNA-seq participants.**

A. Fungal disease was defined as fungal infections other than superficial cutaneous infections (e.g. tinea pedis)

Abbreviations: BMI, body mass index; CAD, coronary artery disease; COPD, chronic obstructive pulmonary disease; DM, diabetes mellitus; HAART, highly active antiretroviral therapy; HIV, human immunodeficiency virus; HLD, hyperlipidemia; HTN, hypertension; PPD, purified protein derivative; PVD, peripheral vascular disease; VL, viral load
